# Neuromodulation of Swarming Behavior in *C. elegans*: Insights into the Conserved role of Calsyntenins

**DOI:** 10.1101/2025.07.06.663344

**Authors:** Navneet Shahi, Nisha Kumari, Sharveri Khapre, Dimple Dahiya, Egemen Saritekin, Aşkın Kocabaş, Babu Kavita

**Affiliations:** Indian Institute of Science, Bangalore, Karnataka, India; Indian Institute of Science Education and Research, Berhampur, India; Vellore Institute of Technology, Vellore, India; Koç University, Istanbul, Türkiye

**Keywords:** Swarming behavior, CASY-1, Serotonin, PDFR-1, Mechanosensation

## Abstract

Collective animal behaviors arise from a complex interplay between internal physiological states and external environmental cues. In *Caenorhabditis elegans*, favorable conditions promote dispersal, while stressors like food scarcity or overcrowding trigger aggregation. Here, we describe a distinct behavior termed as swarming, where *C. elegans* move and feed in aggregates despite abundant food availability. While environmental factors have been implicated in this behavior, the underlying genetic and molecular mechanisms remain unclear. We identify a novel role for the conserved calsyntenin protein CASY-1 in regulating swarming. Through genetic, behavioral, and optogenetic approaches, we show that CASY-1 functions in sensory neurons to modulate the neuropeptide pigment-dispersing factor (PDF) signaling. Mutants in *casy-1* show impaired PDF-1 signaling and reduced inhibition of the serotonin pathway, a known regulator of social behaviors. This dysregulation, along with its associated mechanosensory and foraging defects, likely contributes to the swarming phenotype. Our findings reveal a putative neuromodulatory pathway critical for swarming behavior in *C. elegans*.

## Introduction

The ability of animals to group or disperse is rarely random. It reflects a complex integration of diverse sensory, physiological, and environmental cues (*1-6*). Across species, collective behaviors such as coordinated migration, social feeding, and cooperative defence have evolved as adaptive strategies, seen in ant trails, bird flocks, fish shoals, and mammalian herds (*6-11*). Yet, the underlying neural mechanisms that translate complex cues into coordinated group actions remain poorly understood.

The nematode *Caenorhabditis elegans* offers a powerful system to dissect the genetic and neural basis of collective behavior, owing to its well-characterized nervous system and ease of culturing large populations (*12*). Among its social behaviors, aggregation has been widely studied (*7, 13-15*). While *aggregation* refers to stationary clustering, and *social feeding* or *swarming* describe motile group behaviors, these terms are often used interchangeably in the literature. Here, we define swarming as a distinct, self-organized behavior characterized by the persistent and directed collective movement of *C. elegans* clusters along the food boundary, without substantial dispersal. While environmental factors such as food distribution (*16, 17*), pheromones (*5, 18*), oxygen (*19-21*), pathogen exposure (*21*), and population density (*21*) influence this behavior, its internal molecular and neuromodulatory regulators remains largely unexplored.

Disruptions in social behavior are a hallmark of several neurodevelopmental disorders, including autism, bipolar disorder, and schizophrenia (*7, 22-24*). These impairments are often associated with genes involved in sensory integration and neuromodulation (*6, 7*). Thus, identifying such genes can offer valuable insight into the biological mechanisms underlying social dysfunction in these disorders. In this context, we investigate the role of CASY-1, the *C. elegans* ortholog of mammalian calsyntenin proteins (*25*). Calsyntenins have been linked to multiple neuropsychiatric conditions marked by social deficits (*26-30*). Although CASY-1 has been studied in the context of synaptic transmission (*31-33*), chemotaxis (*25*), and memory formation (*34, 35*), its function in social behavior remains unknown.

Here, we report a novel role for CASY-1 in regulating a distinct collective behavior known as swarming. *casy-1* mutants form stable aggregates at the edge of a bacterial food lawn and fail to disperse, despite abundant food availability. These animals experience starvation and delayed development but continue to cluster, exhibiting a phenotype that resembles compulsive group-seeking behavior (*36*).

Classically, neuropeptide receptor NPR-1 has been widely studied for its role in collective behaviors in *C. elegans* (*19, 37, 38*). However, our findings uncover an additional neuromodulatory circuit involving Serotonin and the Pigment Dispersing Factor-1 (PDF-1). Serotonin is a conserved regulator of social behavior across species. It modulates group level aggregation in locusts (*3*), and shapes context-dependent social responses in crustaceans, fish, rodents, and primates (*39, 40*). Aberrant serotonergic signaling is linked to conditions like autism and social anxiety (*41-43*). In *C. elegans*, serotonin and PDF-1 function antagonistically (*44*). Serotonin inhibits PDF-1 signaling, which in turn regulates arousal (*45, 46*), mating interactions (*47*), and foraging behavior (*46, 48, 49*). We find that *casy-1* mutants show reduced PDF-1 signaling, leading to disinhibition of serotonin pathways and the emergence of a hypersocial swarming phenotype. Through genetic, behavioral, and optogenetic approaches, we show that CASY-1 acts as a key upstream modulator that maintains balance between serotonergic and PDF-1 pathways.

Together, our study defines a previously uncharacterized regulatory mechanism through which CASY-1 constrains collective behavior. This study sheds light on how conserved genetic and neuromodulatory mechanisms shape social organization.

## Materials and Methods

### *C. elegans* strain maintenance

*C. elegans* strains were maintained on nematode growth medium (NGM) agar plates seeded with *E. coli* OP50 at 22°C, following standard protocols (*50*). The Bristol N2 strain served as the wild-type (WT) control. Stage-synchronised populations were obtained using sodium hypochlorite bleaching, as described previously (*51*). Unless otherwise specified, all experiments were conducted using young adult hermaphrodites. A complete list of strains used in this study is provided in Table S2.

### Generation of constructs and Transgenic lines

All plasmids were constructed either using Gibson assembly or restriction cloning and verified by sequencing prior to use. The transcriptional reporter *Pcasy-1::mCherry* was generated by promoter insertion upstream of mCherry in the pPD49.26 vector using restriction digestion. Rescue constructs for each *casy-1* isoform were generated by cloning the respective cDNA downstream of its promoter, with sequences obtained from WormBase as described previously (*32*). *BlaC* constructs were subcloned into the pSF154 Vector (*44*) by adding *Pcasy-1* and *Ppdf-1* promoter sequences. Transgenic *C. elegans* lines were generated by microinjection into the gonads of young adult hermaphrodites, using pCFJ90-mCherry or pCFJ90-GFP as co-injection markers as previously described (*52*). At least three independent lines were generated for each construct; the line with the highest expression penetrance and consistent behavior was selected for analysis. For all the rescue experiments, fluorescent transgenic *C. elegans* were enriched before starting the assay. Plasmids and transgenic strains used in this study are listed in Tables S1-S2, Primers in Table S3.

### CRISPR–Cas9 Genome Editing

CRISPR–Cas9 genome editing was performed following the protocol described in (*53*). Full-gene and domain-specific knockouts of *casy-1* were generated in wild-type *C. elegans*. crRNAs were designed using the CRISPOR tool (*54*), selecting guides with high on-target scores and minimal off-target risk (Table S3). crRNAs, tracrRNAs, and Cas9 protein were obtained from Integrated DNA Technologies (IDT). For N-terminal and C-terminal knockouts, a single-stranded DNA (ssDNA) repair template was also co-injected (Table S3). The injection mix contained 0.25 µg/µl Cas9, 2.5 µM each of gRNA, and TE buffer (pH 7.5), incubated at 37 °C for 15 minutes. The co-injection marker pCFJ90 (5 ng/µl) was added, and the volume was adjusted to 20 µl with nuclease-free water. This mix was microinjected into young adult hermaphrodites. F1 animals expressing the pharyngeal marker were singled out and allowed to produce progeny. Genotyping of parent animals was performed using primers listed in Table S3 (Figures S1 and S2). Mutants carrying the desired deletion were homozygozed and outcrossed three times to WT to remove background mutations.

### Swarming Behavior Recording

Approximately 700–800 synchronized young adult *C. elegans* were collected from 90 mm NGM plates using M9 buffer and washed to remove residual OP50 bacteria. The *C. elegans* were then transferred a few centimetres away from the food lawn on a 60 mm NGM plate seeded with 300 µl E. coli OP50 (OD = 3.0). As WT *C. elegans* also swarm at low food density, we selected a food OD threshold just above this range to ensure that swarming observed was specific to *casy-1* mutants. *C. elegans* behavior was recorded for 10–12 hours at 22 °C under low-light conditions using the MBF Bioscience WormLab tracking system, until the food was completely finished.

### Grayscale Quantification of Swarming

Video recordings were first time-lapsed (∼512×) using OpenShot v3.2.1 software, generating final clips of ∼90 seconds. These were converted to .avi format via VLC media player (3.0.21) for compatibility with FiJi (2.9.0). In FiJi, videos were opened as virtual stacks and converted to 8-bit grayscale images. A Z-projection was created using the “average intensity” method. The “Image Calculator” was then used to subtract the Z-projection from the original video stack, eliminating background features. Binary images of *C. elegans* clusters were generated to remove intensity-based variation, followed by noise filtering using a uniform threshold (2–4 pixels). Mean grayscale intensity was calculated from the processed five equidistant frames and used to quantify swarming behavior. For consistency, only the grayscale intensity value from the mid-point of each video was plotted to compare swarming behavior across genotypes.

### Oxygen-dependent motility measurement

To investigate the oxygen-dependent motility of *C. elegans*, we used a plexiglass-based transparent imaging chamber as described in (*21*). Oxygen concentration ([O₂]) was measured using a standard oxygen sensor (PreSens, Microx TX3). During the [O₂] measurements, the Fiber-optic sensor probe was inserted into the chamber while maintaining a sealed environment. A 50 sccm gas mixture of O₂ and N₂ was used, with flow rates controlled by a flow controller. The oxygen level in the chamber was sequentially reduced from 21% to 0% at 15-minute intervals. Experiments were recorded using a Thorlabs DCC1545M CMOS camera equipped with a Navitar 7000 TV zoom lens. A custom-developed multi-worm tracker code was used to measure the animals’ speed corresponding to specific oxygen levels.

### Single-Animal Foraging Assay

Foraging distance was measured on off-food 90 mm NGM plates lacking cholesterol. Plates were dried to remove excess moisture and equilibrated to room temperature (∼22 °C) prior to use. Well-fed young adult *C. elegans* were gently transferred using an eyelash pick with a drop of halocarbon oil to an intermediate off-food plate to remove residual bacteria, and then placed on the assay plate. The *C. elegans* were allowed to acclimate for 1 minute before recording. Locomotion was recorded for 5 minutes using the MBF Bioscience WormLab imaging system at 8 fps. At least 20 *C. elegans* were recorded per genotype. Animals that moved out of the camera’s field of view during the recording period were excluded from the analysis. Total distance (from the center-point) travelled per animal, as computed by WormLab software, was used as the primary readout and plotted using GraphPad Prism 9.5.1.

### Aggregation Assay

Aggregation was assessed following established protocols (*14, 15*). Fresh 60 mm NGM plates were seeded with 200 µl of *E. coli* OP50 (OD₆₀₀ ∼1.5) to form a ∼2.5-3 cm diameter circular lawn and dried to prevent the formation of a coffee ring. Approximately 70 synchronized young adult *C. elegans* were gently transferred to the center of the food lawn and allowed to settle for 1 hour. Aggregation was manually scored thrice at 15-minute intervals by counting the number of solitary, freely dispersing *C. elegans* and subtracting this from the total number of animals added on the plate. The remaining animals, found in groups near or at the food edge, were considered part of the aggregate. The assay was repeated three times per genotype across three independent days.

### Touch-response Assay

Mechanosensory responses were assessed as previously described (*55*), using 60 mm NGM plates freshly seeded with OP50 (OD ∼1.0). allowed to acclimate for two minutes before testing. All assays were performed blind to genotype. For the nose touch assay, each *C. elegans* was gently stimulated on the nose using an eyelash hair pick, ten times in succession, with a reversal movement scored as a positive response. For the anterior and posterior touch assays, animals were gently touched five times either just behind the pharynx (anterior) or near the tail (posterior). A reversal or forward movement, respectively, was counted as a positive response. Each assay was conducted in four independent replicates, with ten *C. elegans* per replicate. The total number of responses was recorded per animals, and results are reported as the mean percentage of positive responses per animal per genotype.

### Serotonin Supplementation Assay

A 1 M serotonin hydrochloride stock solution (Sigma-Aldrich, SLCQ5039-H9523) was freshly prepared in sterile M9 buffer on the day of the experiment. Working concentrations of 3 mM, 6 mM, and 10 mM were tested based on prior studies (*56*), with 6 mM yielding the most consistent behavioral response. To ensure efficient uptake and minimize degradation during this longer, population-based assay (∼500 *C. elegans* for this assay), animals were pre-exposed to serotonin (100µl per 60mm plate) in their synchronization plates for approximately 2.5 hours before behavioral recording. In parallel, 60 mm NGM assay plates seeded with *E. coli* OP50 were evenly coated with 100 µl of the corresponding serotonin solution and dried in the dark. All plate preparations were conducted under low-light conditions to minimize photodegradation. *C. elegans* were then washed and transferred to one side of the assay plate, as described for swarming assays. Control plates were treated with sterile M9 buffer, and the animals were similarly exposed to buffer-only conditions to control for solvent effects. The data presented correspond to the 6 mM serotonin condition.

### Optogenetics Assay

Optogenetic stimulation was performed using blue-light activation of adenylyl cyclase (BlaC), expressed under a cell-specific promoter as described previously (*44*). Transgenic *C. elegans* were generated by microinjecting the BlaC plasmid into the relevant mutant background, followed by crossing into wild-type (WT) to obtain a consistent genetic background. To prevent unintended activation, transgenic animals were maintained in light-protected containers wrapped in aluminium foil and handled under low-light conditions. Fluorescent animals were enriched before the assay by selecting for co-injection marker expression. For behavioral recordings, the transgenic animals were placed at one end of the bacterial lawn on a 60 mm plate and allowed to reach ∼25% of the food lawn over four hours in the dark. Behavior was then recorded for two hours using the MBF Bioscience WormLab tracking system. The first hour served as a baseline (no stimulation), followed by blue-light exposure during the second hour, delivered in one-minute pulses at six-minute intervals (Figure 4C). The plotted data represent the average grayscale intensity before and after stimulation, with the difference used to quantify the effect of neuronal activation on swarming behavior.

### Statistical analysis

All statistical analyses were performed using GraphPad Prism (9.5.1). The choice of statistical test and graph type depended on the data distribution and experimental design. Data were primarily represented using bar graphs and box-and-whisker plots. Comparisons between two groups were conducted using unpaired two-tailed Student’s t-tests with Welch’s correction. For comparisons across more than two groups, one-way or two-way ANOVA followed by Tukey’s multiple comparison test was applied. Time-course data were plotted as line graphs to visualize phenotypic changes over time. N refers to the number of technical replicates unless otherwise stated. Graphs display mean ± SEM, A p-value < 0.05 was considered statistically significant. For most figures, (*P < 0.05, **P < 0.01, ***P < 0.001, ****P < 0.0001; ns = not significant). Quantification was done using data from at least three independent biological replicates. The specific statistical test used for each dataset is detailed in the corresponding figure legend.

## Results

### CASY-1 regulates swarming via its C-terminal domain in sensory neurons

Our interest into CASY-1’s role in collective behavior began with a serendipitous observation where null mutants of *casy-1* exhibited robust swarm formations. This phenotype emerged spontaneously and reproducibly, prompting us to investigate further.

To quantify swarming, synchronized *C. elegans* were placed at one edge of a uniform bacterial lawn and recorded over 10–12 hours. Time-lapse imaging at five equidistant time points revealed that while WT animals dispersed evenly across the lawn, *casy-1(ind107)* mutants formed stable aggregates at the food boundary. Expression of CASY-1A under its endogenous promoter significantly restored dispersal, although some variability was seen due to mosaic expression from extrachromosomal arrays (Figure 1A).

**Figure 1.**
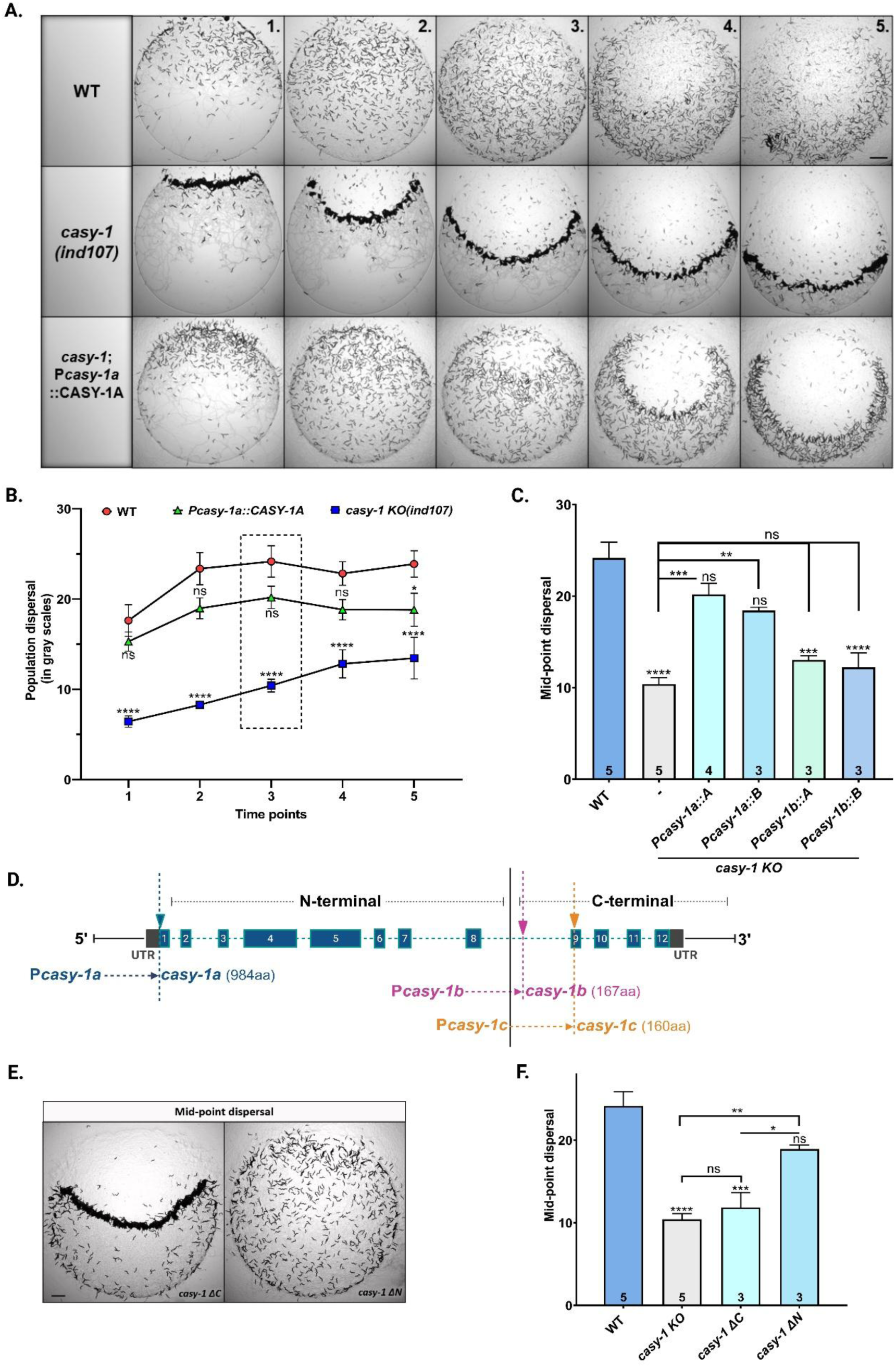
CASY-1 regulates swarming through its C-terminal domain in sensory neurons. (A) Time-course images showing the dispersal dynamics of wild-type (WT), *casy-1* knockout (KO), and endogenous rescue (*Pcasy-1a*::CASY-1A) *C. elegans* across a bacterial lawn at five equidistant time points. (B) Quantification of population dispersal using grayscale intensity across time. Lower grayscale values indicate increased clustering and higher values indicate more dispersal. (C) Mid-point grayscale values used for comparison in *casy-1* mutants and isoform-specific rescue lines. Expression of CASY-1A or CASY-1B under the *Pcasy-1a* promoter significantly rescues the swarming phenotype, whereas their expression under *Pcasy-1b* does not. (D) Schematic of the *casy-1* locus showing three isoforms: CASY-1A (full-length, 984 aa) under *Pcasy-1a* promoter (blue), CASY-1B (167 aa) under *Pcasy-1b* (pink), and CASY-1C (160 aa) under *Pcasy-1c* (orange). Isoforms B and C originate from alternative promoters within eighth intron (∼4000 bp) of the *casy-1a* locus. Transmembrane domain (black) demarcates the N-terminal and C-terminal regions. (E) Mid-point dispersal images of *casy-1* ΔC and *casy-1* ΔN deletions. (F) Quantification of dispersal in WT, *casy-1 KO*, and domain-deleted mutants. Data represent mean ± standard error mean (SEM). N = 3-5 biological replicates, indicated inside the bar graph. Statistical significance was assessed using ordinary one-way ANOVA and Tukey’s multiple comparison test. (*** *P* < 0.001, **** *P* < 0.0001; ns = non-significant). Scale bar = 20 mm.

Occasional WT aggregation was seen at early or late time points (frames 1-2 or 4-5), typically due to edge entry or food depletion, but this was transient and less prominent than *casy-1* mutants. The assay midpoint was selected for consistent quantification across replicates. Swarming was analyzed using a grayscale-based method, where higher values indicate more dispersal (Figure 1B). Mid-point values were compared across genotypes (Figure 1C).

The *casy-1* gene encodes three isoforms through alternative promoter usage (Figure 1D). The full-length CASY-1A, containing all conserved domains, is expressed predominantly in sensory and interneurons. Shorter isoforms CASY-1B and CASY-1C lack the N-terminal region and are expressed in ventral motor neurons. To determine whether CASY-1 regulates swarming through its function in sensory or motor neurons, we performed isoform-specific rescue experiments using the *Pcasy-1a* and *Pcasy-1b* promoters. Due to sequence similarity and overlapping expression, *Pcasy-1b* and *Pcasy-1c* are often used interchangeably (*32*).

Expression of either CASY-1A or CASY-1B under *Pcasy-1a* significantly rescued the swarming phenotype. In contrast, motor neuron–restricted expression under *Pcasy-1b* failed to do so (Figure 1C). This suggests that CASY-1 acts in sensory neurons and that the C-terminal region alone is sufficient.

To test this directly, we generated CRISPR-based *casy-1* full and domain deletions (Figure S1A-S1B). Deletion of the C-terminal domain (*ΔC*) fully phenocopied the *casy-1* null mutants. In contrast, deletion of the N-terminal domain (*ΔN*) had little or no effect (Figures 1E–F). These findings establish that the C-terminal domain is necessary and sufficient for regulating swarming, likely via sensory neurons.

One experimental consideration while maintaining *casy-1* mutants is their tendency to remain aggregated and failure to disperse, which complicates synchronization and uniform growth. Despite bleaching, these mutants often develop asynchronously, likely due to uneven food access and starvation. To minimize this, we plated fewer eggs at opposite ends of larger 90 mm NGM plates to improve dispersal during early development. Although slight variation in developmental stage persisted, population size was carefully calibrated across replicates to ensure comparability.

To validate the robustness of the phenotype, we compared the CRISPR knockout line *casy-1(ind107)* that we had generated with two previously reported alleles, *tm718* and *wp78* (Illustrated in Figure S1C). All three alleles exhibited comparable swarming behaviors (Figure S1D). Figure S1A shows the genomic deletion in the *casy-1(ind107)* line and its genotyping. We also reproduced previously reported Aldicarb and PTZ sensitivity phenotypes associated with the *tm718* mutant line (*32, 57*) in the *casy-1(ind107)* animals (Figures S1E-F). All subsequent experiments were performed with the *casy-1(ind107)* allele.

### Swarming in *casy-1* mutants is independent of known extrinsic sensory and chemical cues

To test whether extrinsic cues contribute to the swarming seen in *casy-1* mutants, we first examined the neuropeptide receptor NPR-1, which is a known regulator of social behaviors like aggregation, social feeding, and swarming (*14, 17, 37, 38, 58*). In *npr-1* mutants, these phenotypes are partly driven by altered oxygen sensing (*19-21*). Our previous work also showed that *casy-1* interacts genetically with *npr-1* to regulate synaptic signaling (*33*). This prompted us to explore whether *casy-1* and *npr-1* influence swarming through similar mechanisms.

First, we compared aggregation patterns across genotypes. As expected, *npr-1* mutants formed multiple robust clusters along the food border. In contrast, *casy-1* animals remained relatively dispersed, resembling WT animals (Figure 2A). This indicates that classical aggregation in *npr-1* mutants is not necessary for swarm formation in *casy-1* mutants. Next, we compared swarming patterns across genotypes. Unlike *npr-1* mutants, which formed dispersed swarms emerging from food-depleted zones, *casy-1* mutants formed compact and localized swarms. Interestingly, *casy-1; npr-1* double mutants displayed a composite phenotype, characterized by the formation of swarm-like clusters similar to *casy-1* mutants, along with enhanced peripheral dispersal seen in *npr-1* mutants (Figure 2B). Although the overall dispersal area was comparable among *casy-1, npr-1,* and double mutants, the spatial organization and dynamic distribution on the food lawn differed markedly (Figures 2B-2C), suggesting that *casy-1* and *npr-1* regulate distinct facets of swarming behavior.

**Figure 2.**
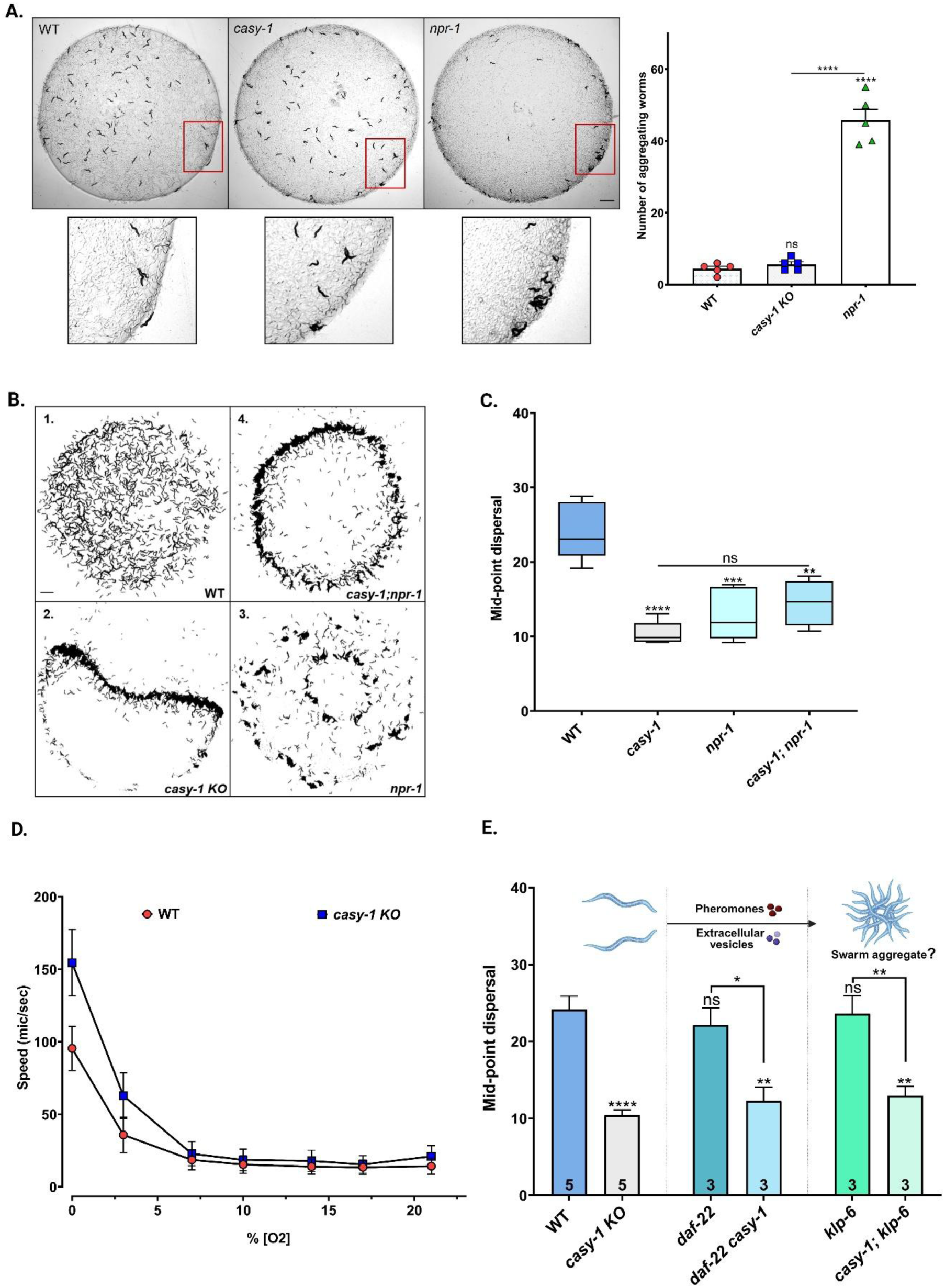
*casy-1* mutants exhibit a distinct swarming phenotype from *npr-1* and the swarming is not driven by extrinsic factors. (A) Representative images showing aggregation patterns in WT, *casy-1*, and *npr-1* mutants on a food lawn. Zoomed insets highlight *C. elegans* clustering at the food border. Quantification of the number of aggregated *C. elegans* indicates that *casy-1* mutants resemble WT and do not exhibit the border aggregation phenotype characteristic of *npr-1* mutants. (B) Swarming assay snapshots at the mid-point showing distinct dispersal patterns in 1. WT, 2. *casy-1*, 3. *npr-1*, and 4. *casy-1; npr-1* double mutants. (C) Quantification of mid-point dispersal across genotypes based on grayscale intensity measurements. The box-and-whisker plot displays the median and data range. While *casy-1* and *npr-1* mutants exhibit reduced dispersal compared to WT, no significant difference is observed between these single and double mutants. (D) Oxygen response assay comparing locomotor speed of *casy-1* and WT animals across a range of oxygen concentrations. No significant difference was observed, indicating oxygen sensitivity is not altered in *casy-1* mutants. (E) Quantification of relative dispersal in mutants defective in pheromone synthesis (*daf-22*) and extracellular vesicle biogenesis (*klp-6*), as well as their double mutants with *casy-1* (as *daf-22* and *casy-1* express on the same chromosome, they are denoted *daf-22 casy-1* unlike *casy-1; klp-6).* Neither pathway significantly alters the swarming phenotype, suggesting that the swarming behavior is independent of these extrinsic communication mechanisms. Data represent mean ± SEM. Biological replicates are indicated within each bar. Statistical analysis was performed using one-way ANOVA followed by Tukey’s multiple comparison test (* *P* < 0.05, ** *P* < 0.01, *** *P* < 0.001, **** *P* < 0.0001; ns = non-significant). Scale bar = 20 mm.

To assess whether locomotory speed might explain these differences, we measured individual *C. elegans* velocities. Mutants in *npr-1* exhibit significantly higher speeds then WT animals as shown previously ((*14*) and Figure S2A). The *npr-1* mutants also showed increased speed when compared to *casy-1*, consistent with their broader dispersal (Figure S2A). However, *C. elegans* speed alone could not fully explain the behavior as *casy-1; npr-1* double mutants despite having WT-like speed still exhibited swarm formation (Figure 2C). It is worth noting that these are single animal speed measurements and may not indicate how speed changes in a swarm. We next asked whether *casy-1* mutants display altered oxygen sensitivity, as seen in *npr-1* (*19, 21*). Unlike *npr-1* mutants, *casy-1* animals responded similarly to WT across varying oxygen concentrations (Figure 2D), ruling out hypoxia-driven clustering. Swarming in WT animals typically occurs only at high population densities (*21*), while *npr-1* mutants can form small swarms by sensing local oxygen depletion. *casy-1* mutants also show density-dependent clustering, but their swarming arises through a mechanism independent of oxygen sensing. Further, we asked whether defective food sensation contributes to the phenotype. *odr-3* mutants, which are impaired in olfactory food detection, dispersed normally like WT animals (Figure S2B). This suggests that food perception is not responsible for the swarming seen in *casy-1* mutants.

When *C. elegans* encounter one another in close proximity, they can engage in short-range communication that modulates social behaviors, including aggregation. Two key mediators of such inter-animal signaling are pheromones and extracellular vesicles (EVs), both of which have been implicated in regulating social interactions (*15, 18*). To test whether these cues contribute to the swarm formation in *casy-1* mutants, we examined *daf-22* and *klp-6* mutants. *daf-22* is required for the biosynthesis of ascaroside pheromones (*15, 18*), while *klp-6* is involved in EV release from sensory neurons (*59*). Both single mutants exhibited WT-like dispersal, and the *daf-22 casy-1* and *klp-6; casy-1* double mutants continued to swarm like *casy-1* alone (Figure 2E). These results rule out a role for pheromone or EV-based communication in driving swarming in *casy-1* mutants.

Taken together, our findings indicate that the swarming behavior in *casy-1* mutants is not caused by altered oxygen sensing, food perception, pheromone signaling, or EV-mediated cues. Instead, it is more likely driven by changes in intrinsic pathways and the regulation of locomotor states.

### Elevated serotonin signaling underlies swarming behavior in *casy-1* mutants

Having ruled out known extrinsic regulators of collective behavior, we asked whether intrinsic neuromodulatory pathways contribute to the *casy-1* swarming phenotype. Prior work has linked CASY-1’s C-terminal domain to the vesicular trafficking and release of neuromodulators (*32, 33, 60*), prompting us to assess whether neurotransmitter imbalance underlies this behavior.

To investigate this, we analyzed a panel of neurotransmitter-deficient mutants, including those affecting glutamate, GABA, acetylcholine, dopamine, and serotonin pathways. None of these mutants displayed swarming behaviors, suggesting that the *casy-1* phenotype is not due to loss of canonical neurotransmission (Figures S3A–B). Given serotonin’s conserved role in social behaviors across species species (*3, 39, 40, 43, 61*), and its involvement in modulating *C. elegans* foraging (*44, 62, 63*), we focused on this pathway.

To examine whether serotonin contributes to swarming, we supplemented WT animals with exogenous serotonin. This treatment was sufficient to induce swarming (Figure 3A), although the response was weaker than that seen in untreated *casy-1* mutants. In contrast, *casy-1* mutants were unaffected by serotonin (Figure 3A-B), suggesting that their serotonergic signaling might already be above baseline. To test whether this was a general effect of biogenic amines, we supplemented *C. elegans* with dopamine. Despite literature supporting dopamine’s role in social and aggregative behaviors (*64, 65*), neither WT nor *casy-1* animals displayed any change in dispersal (Figures S3C -D).

**Figure 3.**
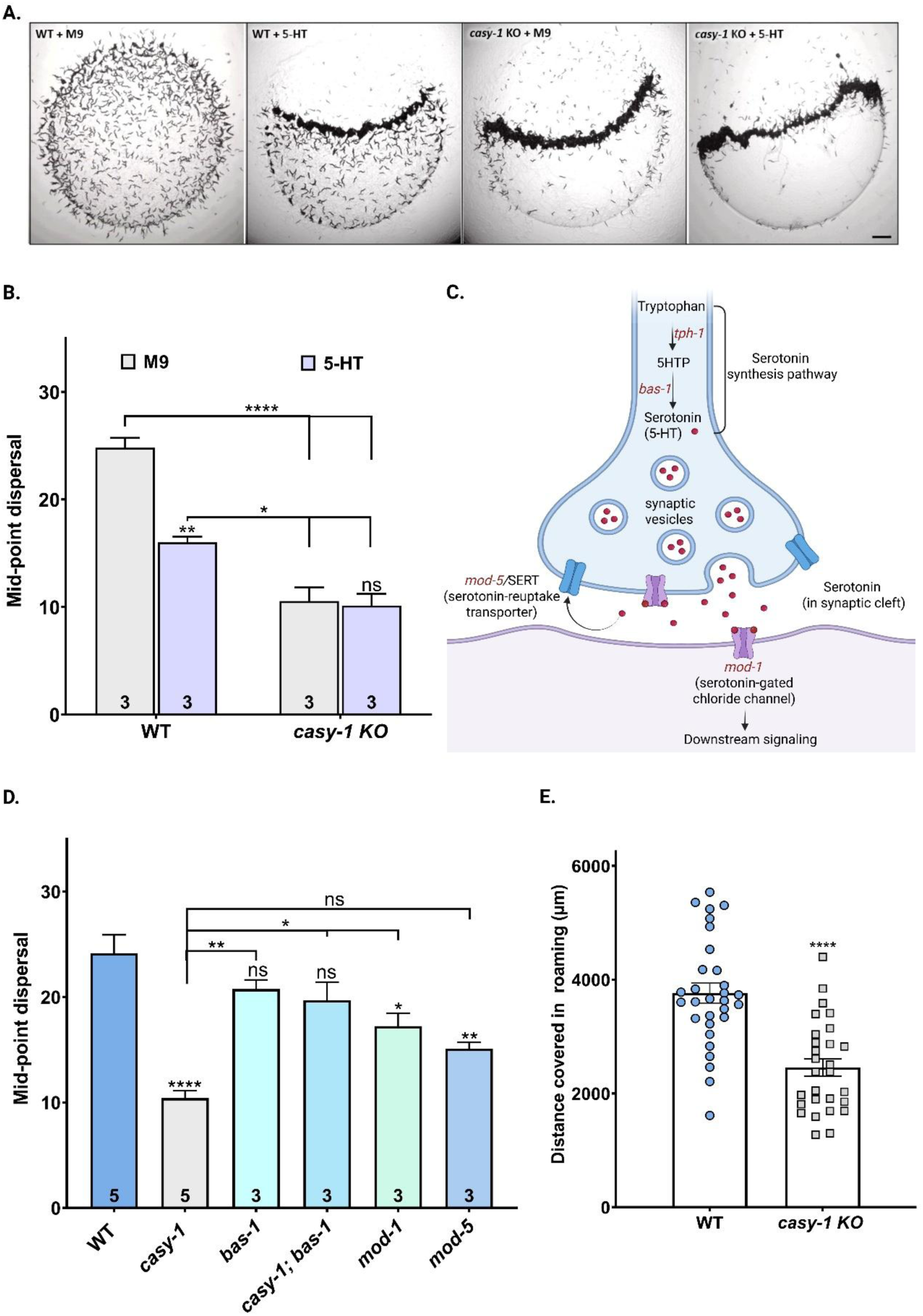
Serotonin signaling contributes to swarming behavior in *casy-1* mutants. (A) Images showing the effect of serotonin (5-HT) supplementation on WT and *casy-1* mutant animals, with M9 buffer as a control (B) Relative dispersal at the mid-point for WT and *casy-1* mutants supplemented with serotonin (5-HT, purple) or solvent control (M9, grey). WT animals show reduced dispersal with serotonin, while *casy-1* mutants exhibit no change. (C) Diagram of the serotonin signaling pathway. Tryptophan is converted to 5-HTP by *tph-1*, and then to serotonin by *bas-1*. Serotonin is packaged into vesicles, released into the synaptic cleft, and signals via *mod-1*, a serotonin-gated chloride channel. *mod-5* encodes the serotonin reuptake transporter (SERT). Image created with BioRender. (D) Relative dispersal at the mid-point for the indicated genotypes. Serotonin-deficient mutants (*bas-1*, *casy-1; bas-1*) resemble WT, while mutants with elevated serotonin signaling (*mod-1*, and *mod-5*) display reduced dispersal similar to *casy-1*. (E) Scatter plot showing total distance covered during roaming by individual WT and *casy-1* animals. *casy-1* mutants exhibit significantly reduced movement. Data are presented as mean ± SEM. For B and D, n = 3 biological replicates (plates) per group; replicate numbers are indicated within bars in panel D. For E, ∼30 individual animals per genotype were analyzed. Statistical analysis was performed using one-way ANOVA with Tukey’s post hoc test for B and D, and unpaired *t*-test with Welch’s correction for E. Significance: (* *P* < 0.05, ** *P* < 0.01, *** *P* < 0.001, **** *P* < 0.0001; ns = non-significant). Scale bar = 20 mm.

To further dissect serotonin’s role, we examined its signaling pathway mutants (Figure 3C). Mutants deficient in serotonin synthesis (*tph-1* and *bas-1*) showed WT dispersal (Figures S3A, 3D). We couldn’t obtain double mutants of *casy-1* and *tph-1* as they are located close on the second chromosome. However, the *casy-1; bas-1* double mutants exhibited a strong suppression of the swarming observed in *casy-1* single mutants, reinforcing serotonin excess as a causal factor (Figure 3D). We next examined *mod-1* mutants, lacking the inhibitory serotonin-gated chloride channel. These animals exhibited moderate aggregation, though less pronounced than in *casy-1*. Interestingly, although *mod-1* mutants have been associated with increased exploration (*44, 62, 63*), we observed a mild swarming phenotype. In contrast, *mod-5* mutants, which accumulate extracellular serotonin due to defective reuptake, formed stable swarms similar to *casy-1* (Figure 3D).

To further understand the behavioral effect of elevated serotonin, we quantified exploration in *casy-1* mutants. These animals showed significantly reduced exploratory behavior compared to WT (Figure 3E). In *C. elegans*, serotonin typically promotes a shift from roaming to dwelling in response to food availability (*44, 62, 63, 66*). However, the clustering we observe in *casy-1* mutants differs from classic dwelling. These animals remain at the food boundary, rather than settling within the lawn, and show slowed locomotion even before food contact (Figure S3A). This suggests that the behavioral state is not triggered by food ingestion, but instead reflects a serotonin-dependent internal state that mimics features of dwelling. Together, these findings support a model in which elevated serotonergic signaling contributes to the swarm formation in *casy-1* mutants.

### PDF-1 signaling and mechanosensation act downstream of CASY-1 to regulate swarming

The behavioral features of *casy-1* mutants such as reduced exploration and slower locomotion (Figure S2A, Figure 3E), resemble a dwelling-like state (*44, 49*). Since serotonin and Pigment dispersing factor 1 (PDF-1) function antagonistically to regulate dwelling and roaming in *C. elegans* (Figure 4A). we investigated whether reduced PDF-1 signaling contributes to the swarming observed in *casy-1* mutants.

**Figure 4.**
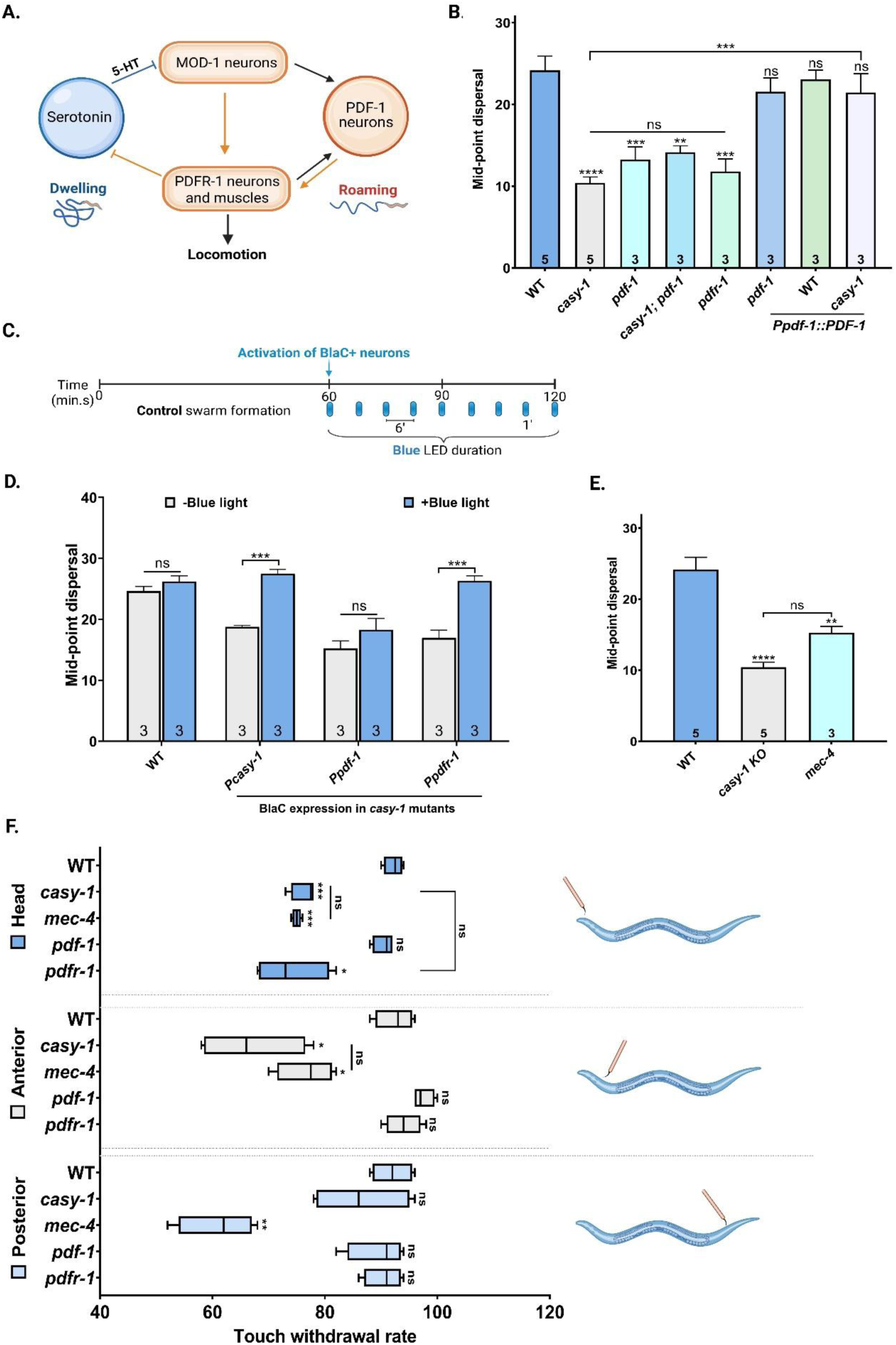
CASY-1 regulates swarming via PDF-1 neuromodulatory balance and mechanosensory control. (A) Schematic of the neuromodulatory circuit regulating behavioral state in C. elegans, highlighting antagonistic interactions between serotonin and PDF-1 pathways. Serotonin (5-HT), produced by NSM/HSN neurons (blue), promotes dwelling by inhibiting MOD-1–expressing neurons. PDF-1 (orange), released from AVB and PVP neurons acts via the receptor PDFR-1 to promote roaming. Adapted from (*63*), created using BioRender. (B) Quantification of mid-point dispersal. *casy-1*, *pdf-1*, *casy-1; pdf-1*, and *pdfr-1* mutants all display reduced dispersal compared to WT, consistent with swarming. Endogenous expression of *pdf-1* rescues dispersal in *pdf-1* mutants and *pdf-1* overexpression in *casy-1* mutants fully suppresses swarming. (C) Schematic showing the optogenetic stimulation paradigm using blue-light illumination (using Biorender). (D) Quantification of dispersal before and after blue-light stimulation in strains expressing BLaC under different promoters. Blue light had no effect on WT controls (expressing *Pcasy-1::BLaC*). In contrast, stimulation of *Pcasy-1–* or *Ppdfr-1–*driven BLaC in *casy-1* mutants significantly increased dispersal, while *Ppdf-1* activation showed no effect. (E) Mid-point dispersal quantification in *casy-1* and *mec-4* mutants. *mec-4* animals, defective in gentle touch sensation, show reduced dispersal similar to *casy-1*. (F) Gentle touch responses scored as withdrawal rate (%) upon stimulation at head, anterior, and posterior body regions. *casy-1* and *pdfr-1* mutants show impaired head touch response, with *casy-1* also showing mild anterior defects. *mec-4* mutants serve as a negative control. Bar plots in B, D, and E represent mean ± SEM. Touch assays (F) are shown as box plots across four biological replicates (N = 10 *C. elegans* each). Statistical tests: one-way ANOVA with Tukey’s post hoc test (B, D, E); two-way ANOVA with Tukey’s test (F). (* *P* < 0.05, *** *P* < 0.001, **** *P* < 0.0001; ns = non-significant).

Mutants for *pdf-1*, *pdfr-1*, and *casy-1; pdf-1* double mutants showed persistent swarming, similar to *casy-1* alone (Figure 4B). Expression of PDF-1 under its endogenous promoter rescued swarming in *pdf-1* mutants, confirming PDF-1’s role. Interestingly, overexpression of PDF-1 in *casy-1* mutants suppressed swarming, suggesting that PDF-1 signaling is compromised in the absence of CASY-1 (Figure 4B).

To test circuit involvement, we optogenetically activated neurons expressing either PDF-1 or PDFR-1 (Figure 4C). Strikingly, blue-light stimulation of PDFR-1–expressing neurons fully dispersed swarm aggregates, while activation of PDF-1–expressing neurons produced little to no effect (Figure 4D). One possible explanation for this could be that *casy-1* mutants have impaired PDF-1 trafficking, limiting its release and function, despite optogenetic stimulation. In contrast, direct activation of PDFR-1–expressing neurons bypasses this defect and restores dispersal, suggesting that PDFR-1 neuron activity is sufficient to suppress swarming and acts downstream of *casy-1*. Further, Control WT animals remained dispersed regardless of light exposure. Notably, light activation of CASY-1–expressing neurons also suppressed swarming, similar to PDFR-1 activation (Figure 4D).

We also assessed whether this was a general defect in neuropeptide signaling. Both *egl-3* (neuropeptide processing) and *unc-31* (neuropeptide release) mutants, as well as their double mutants with *casy-1*, also showed swarming phenotypes (Figure S4A). A screen of other sensory-neuron-expressed neuropeptides and their receptors previously used to study foraging behaviors in our laboratory (*67*), revealed variable phenotypes (Figure S4B), indicating that diverse neuromodulatory pathways can influence swarming.

Next, we examined whether downstream signaling through PDFR-1 contributes to the swarming phenotype. PDFR-1 is expressed in mechanosensory neurons and muscles. It has been linked to touch sensitivity and mating-associated contact sensitivity (*47, 49, 68*). We tested whether touch sensitivity influences swarming by analyzing *mec-4* mutants, which are defective in gentle touch (*69, 70*), exhibited moderate swarming (Figure 4E).

Mechanosensory feedback is known to affect collective behavior in *C. elegans* and other species (*2, 71*). To explore this, we measured head, anterior, and posterior touch responses in *casy-1*, *pdf-1*, and *pdfr-1* mutants, using *mec-4* as a control. *casy-1* mutants showed impaired responses to head and anterior touch. *pdfr-1* mutants showed reduced head sensitivity. *pdf-1* and WT animals responded normally. Posterior touch responses were intact in all genotypes (Figure 4F).

These results suggest that *casy-1* mutants have reduced anterior mechanosensation. This may lower their ability to respond to nearby animals and help maintain clusters. We propose that impaired touch sensation, combined with neuromodulatory imbalance, supports the emergence and stability of swarming in *casy-1* mutants.

### Mechanistic model for CASY-1–mediated neuromodulatory control of swarming in *C. elegans*

This model summarizes the cellular and behavioral changes observed in *casy-1* mutants. Loss of CASY-1 leads to reduced PDF-1/PDFR-1 signaling and a relative increase in serotonin activity. This neuromodulatory imbalance is associated with slower movement, reduced exploration, and impaired mechanosensation. Together, these behavioral shifts likely contribute to the emergence and maintenance of swarming in *casy-1* mutants (Figure 5).

**Figure 5.**
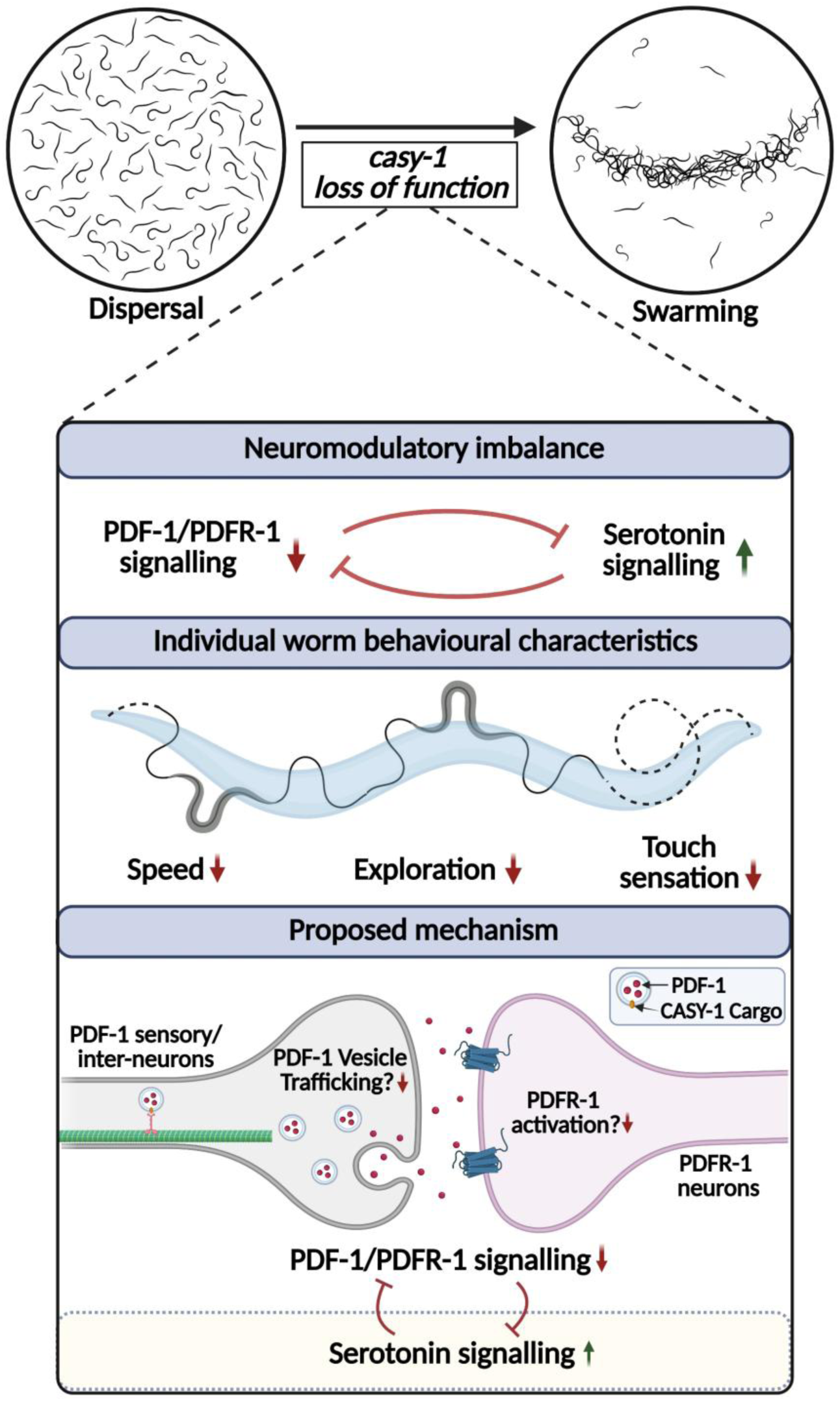
Model for CASY-1–mediated regulation of swarming in *C. elegans.* The schematic illustrates how *casy-1* null mutation shifts animals from a dispersed state to a swarming state. This transition is mediated by a neuromodulatory imbalance between the PDF-1/PDFR-1 pathway (↓) and the serotonergic system (↑). Reduced PDF-1 signaling fails to suppress serotonin activity, resulting in elevated serotonin levels. The imbalance affects individual behavioral traits, including decreased speed, reduced exploration, and impaired touch sensitivity, ultimately driving swarm formation. The lower panel shows a proposed circuit mechanism: CASY-1 functions in PDF-1 expressing neurons, potentially affecting vesicle trafficking. Loss of CASY-1 reduces PDFR-1 activation in downstream neurons, leading to increased serotonergic signaling and altered behavior. The *C. elegans* trail is adapted from (*79*) and all illustrations were created with BioRender.

Previous work has shown that PDF-1 is expressed in sensory neurons such as PVP, AVB, and SIAV, where it modulates roaming and dwelling behaviors via activation of its receptor PDFR-1 on interneurons including AIY, RIM, and RIA (*44*). These interneurons serve as critical hubs for behavioral state transitions, integrating sensory input and internal signals to regulate locomotor dynamics.

CASY-1 is reported to be expressed in several PDF-1 producing sensory neurons (*32, 72, 73*). Further, expression of CASY-1 in in sensory neurons of *casy-1* mutants is sufficient to rescue the swarming phenotype, suggesting that CASY-1 likely acts within this population to regulate PDF-1 signaling. Given that the C-terminal domain is essential for rescue, and has been implicated in vesicular trafficking (*31-33*), we propose that CASY-1 may support the intracellular trafficking or release of PDF-1. This would impact the activation of its receptor, PDFR-1, in downstream interneurons and muscles. In *casy-1* mutants, reduced PDF-1 signaling likely leads to a disinhibition of serotonergic pathways, contributing to elevated serotonin levels. This shift promotes a dwelling-like state and reduced roaming. As a feedback loop, enhanced serotonin may further reinforce aggregation.

Together, this model connects molecular, cellular, and behavioral findings to propose a mechanism by which CASY-1 regulates swarming through modulation of neuromodulatory balance and circuit-level interactions.

## Discussion

Collective behaviors have been extensively examined through ethological and ecological lenses, offering key insights into the rules that guide group organization and dynamics (*11, 74-76*). However, the genetic and molecular mechanisms underlying these behaviors remain relatively unexplored. This gap may stem from the challenge of linking emergent population-level phenomena to specific molecular mechanisms. In this study, we leverage the genetic tractability of *C. elegans* to explore how a conserved synaptic protein, CASY-1, regulates a distinct collective behavior, termed swarming via neuromodulatory signaling.

We identify a self-organized, persistent swarming phenotype in *casy-1* mutants that emerges independently of environmental stressors. Unlike canonical aggregation responses triggered by hypoxia, starvation or crowding (*16, 17, 19-21*), swarming in *casy-1* mutants occurs even in nutrient-rich conditions (Figure 1). This persistent clustering at the food boundary impairs optimal development, highlighting an ecological trade-off associated with excessive social behavior. Notably, swarming appears to be self-reinforcing. For instance, once a small group began swarming, neighboring animals were more likely to join. However, we found no evidence for mediation by pheromones or extracellular vesicles (Figure 2), suggesting the involvement of intrinsic pathways.

Our results reveal a conserved role for neuromodulatory balance in the emergence of swarming. We show that *casy-1* mutants have reduced Pigment-dispersing factor-1 (PDF-1) signaling, which correlates with elevated serotonin activity (Figures 3-5). Serotonin is a conserved regulator of social behavior and has been implicated in disorders such as autism and depression (*3, 39, 40, 43, 61*). In *C. elegans*, serotonin promotes dwelling, while PDF-1 promotes roaming, and the two act antagonistically to regulate behavioral states (*44*). PDF-1 also tunes behavioral states in an evolutionary conserved manner, where in *Drosophila*, PDF controls circadian and arousal behaviors (*77*), and its mammalian homolog, vasoactive intestinal peptide (VIP), integrates neuromodulatory cues with circadian input to control arousal (*78*).

A key mechanistic question is how CASY-1 functions within this circuit. We show that the C-terminal domain of CASY-1 is essential for its function (Figure 1), consistent with its previously reported role in vesicular trafficking (*32, 35, 60*). We did not observe significant changes in the transcript levels of *pdf-1* or its receptor *pdfr-1* (data not shown), suggesting that CASY-1 likely acts at a post-transcriptional level. We propose that CASY-1 facilitates the trafficking or release of PDF-1 neuropeptides (Figure 5). Supporting this, PDF-1 overexpression suppresses swarming in *casy-1* mutants, and optogenetic activation of PDFR-1–expressing neurons disperses swarms (Figure 4). In contrast, activating PDF-1– producing neurons has no effect, suggesting that the signaling deficit lies upstream of receptor activation. Together, these results position CASY-1 upstream of PDF-1 signaling in regulating collective behavior.

While our data support a model in which neuromodulatory imbalance drives swarming, some limitations remain. First, although grayscale-based analysis of population dispersal is useful for distinguishing between clearly swarming and non-swarming genotypes, it is less effective for mutants displaying intermediate or context-dependent behaviors.

Quantitative metrics that better capture the degree and dynamics of aggregation will be useful. Second, we lack a clear understanding of how individual behavioral traits, such as movement characteristics, scale into group-level behavior. Although, *casy-1* single mutants show reduced locomotion and exploration (Figures 2, 3), their transition into clusters may also involve altered local interactions, possibly triggered by proximity to neighboring *C. elegans* in the swarm. Additionally, *casy-1* mutants exhibit impaired anterior mechanosensation, a phenotype shared with *pdfr-1* mutants (Figure 4, (*68*)). This led us to speculate that reduced touch responsiveness may stabilize aggregates once formed. However, how these sensory changes integrate with neuromodulatory states to promote swarming remains unclear.

In summary, this study uncovers a novel role for CASY-1 in regulating swarming behavior through a conserved neuromodulatory pathway. Future work combining circuit-level manipulations with high-resolution behavioral modelling will be essential to dissect the neural mechanisms that shape group dynamics. Such insights could extend to broader contexts, including the neural basis of social behavior and its dysregulation in neurodevelopmental disorders.

## Supporting information

Combined supplemental file

## Acknowledgements

We thank Serena Ding and André Brown for insightful suggestions on experimental design, Rohit Sachdeva and Kamal Kishore for his contributions to video processing and quantification. We acknowledge Hernán Jaramillo and Cori Bargmann for providing the PSF154 plasmid, Kamal Kishore and Jagan J for BAB8047 plasmid. We thank Isabel Beets, Umer Saleem, and Sharanya for generously sharing some strains used in this study. Some strains were obtained from the Caenorhabditis Genetics Center (CGC), supported by the NIH Office of Research Infrastructure Programs (P40 OD010440), and others were provided by the National Bioresource Project (NBRP), Japan. All illustrations were created with BioRender. We sincerely thank Jagan J for help with CRISPR, and Inchara M for assistance with experiments, and Palagiri Suresh for invaluable routine support. We are also grateful to all members of the Kavita Babu lab for their thoughtful suggestions and feedback throughout the course of this work and for critical reading of this manuscript.

## Author Contributions

NS: design and execution of experiments, data analyses, manuscript writing/editing and supervision, NK: execution of experiments and manuscript editing, SK: execution of experiments, DD: execution of experiments, AK and ES performed oxygen-dependent motility measurements, and KB: supervision, funding acquisition, and manuscript writing/editing.

## References

1. I. D. Couzin, Collective animal migration. Curr Biol 28, R976–R980 (2018).

2. P. Ramdya et al., Mechanosensory interactions drive collective behaviour in Drosophila. Nature 519, 233–236 (2015).

3. M. L. Anstey, S. M. Rogers, S. R. Ott, M. Burrows, S. J. Simpson, Serotonin mediates behavioral gregarization underlying swarm formation in desert locusts. Science 323, 627–630 (2009).

4. A. Selimbeyoglu et al., Modulation of prefrontal cortex excitation/inhibition balance rescues social behavior in CNTNAP2-deficient mice. Sci Transl Med 9, (2017).

5. C. S. Muirhead, J. Srinivasan, Small molecule signals mediate social behaviors in C. elegans. J Neurogenet 34, 395–403 (2020).

6. R. Harpaz et al., Collective behavior emerges from genetically controlled simple behavioral motifs in zebrafish. Sci Adv 7, eabi7460 (2021).

7. M. H. Cowen et al., Conserved autism-associated genes tune social feeding behavior in C. elegans. Nat Commun 15, 9301 (2024).

8. I. Janczarek, A. Wisniewska, M. H. Chruszczewski, E. Tkaczyk, A. Gorecka-Bruzda, Social Behaviour of Horses in Response to Vocalisations of Predators. Animals (Basel) 10, (2020).

9. F. Hiramatsu, J. W. Lightfoot, Kin-recognition and predation shape collective behaviors in the cannibalistic nematode Pristionchus pacificus. PLoS Genet 19, e1011056 (2023).

10. M. J. Hansen, P. Domenici, P. Bartashevich, A. Burns, J. Krause, Mechanisms of group-hunting in vertebrates. Biol Rev Camb Philos Soc 98, 1687–1711 (2023).

11. D. J. Sumpter, The principles of collective animal behaviour. Philos Trans R Soc Lond B Biol Sci 361, 5–22 (2006).

12. M. C. Leung et al., Caenorhabditis elegans: an emerging model in biomedical and environmental toxicology. Toxicol Sci 106, 5–28 (2008).

13. H. Jang et al., Dissection of neuronal gap junction circuits that regulate social behavior in Caenorhabditis elegans. Proc Natl Acad Sci U S A 114, E1263–E1272 (2017).

14. M. de Bono, C. I. Bargmann, Natural variation in a neuropeptide Y receptor homolog modifies social behavior and food response in C. elegans. Cell 94, 679–689 (1998).

15. E. Z. Macosko et al., A hub-and-spoke circuit drives pheromone attraction and social behaviour in C. elegans. Nature 458, 1171–1175 (2009).

16. S. S. Ding, L. S. Muhle, A. E. X. Brown, L. J. Schumacher, R. G. Endres, Comparison of solitary and collective foraging strategies of Caenorhabditis elegans in patchy food distributions. Philos Trans R Soc Lond B Biol Sci 375, 20190382 (2020).

17. S. S. Ding, L. J. Schumacher, A. E. Javer, R. G. Endres, A. E. Brown, Shared behavioral mechanisms underlie C. elegans aggregation and swarming. Elife 8, (2019).

18. M. de Bono, D. M. Tobin, M. W. Davis, L. Avery, C. I. Bargmann, Social feeding in Caenorhabditis elegans is induced by neurons that detect aversive stimuli. Nature 419, 899–903 (2002).

19. B. H. Cheung, M. Cohen, C. Rogers, O. Albayram, M. de Bono, Experience-dependent modulation of C. elegans behavior by ambient oxygen. Curr Biol 15, 905–917 (2005).

20. J. M. Gray et al., Oxygen sensation and social feeding mediated by a C. elegans guanylate cyclase homologue. Nature 430, 317–322 (2004).

21. E. Demir, Y. I. Yaman, M. Basaran, A. Kocabas, Dynamics of pattern formation and emergence of swarming in Caenorhabditis elegans. Elife 9, (2020).

22. T. Elmer, C. Stadtfeld, Depressive symptoms are associated with social isolation in face-to-face interaction networks. Sci Rep 10, 1444 (2020).

23. C. A. Depp et al., Social competence and observer-rated social functioning in bipolar disorder. Bipolar Disord 12, 843–850 (2010).

24. M. C. Roozen, M. J. H. Kas, Assessing genetic conservation of human sociability-linked genes in C. elegans. Behav Genet 55, 141–152 (2025).

25. D. D. Ikeda et al., CASY-1, an ortholog of calsyntenins/alcadeins, is essential for learning in Caenorhabditis elegans. Proc Natl Acad Sci U S A 105, 5260–5265 (2008).

26. C. Preuschhof et al., KIBRA and CLSTN2 polymorphisms exert interactive effects on human episodic memory. Neuropsychologia 48, 402–408 (2010).

27. A. Vagnoni et al., Calsyntenin-1 mediates axonal transport of the amyloid precursor protein and regulates Abeta production. Hum Mol Genet 21, 2845–2854 (2012).

28. S. V. Ranneva, K. S. Pavlov, A. V. Gromova, T. G. Amstislavskaya, T. V. Lipina, Features of emotional and social behavioral phenotypes of calsyntenin2 knockout mice. Behavioural brain research 332, 343–354 (2017).

29. K. Mori et al., Loss of calsyntenin paralogs disrupts interneuron stability and mouse behavior. Mol Brain 15, 23 (2022).

30. T. V. Lipina et al., Cognitive Deficits in Calsyntenin-2-deficient Mice Associated with Reduced GABAergic Transmission. Neuropsychopharmacology 41, 802–810 (2016).

31. S. Thapliyal, S. Ravindranath, K. Babu, Regulation of Glutamate Signaling in the Sensorimotor Circuit by CASY-1A/Calsyntenin in Caenorhabditis elegans. Genetics 208, 1553–1564 (2018).

32. S. Thapliyal et al., The C-terminal of CASY-1/Calsyntenin regulates GABAergic synaptic transmission at the Caenorhabditis elegans neuromuscular junction. PLoS Genet 14, e1007263 (2018).

33. N. Shahi, S. Thapliyal, K. Babu, Sensory modulation of neuropeptide signaling by CASY-1 gates cholinergic transmission at Caenorhabditis elegans neuromuscular junction. J Biosci 50, (2025).

34. F. J. Hoerndli et al., A conserved function of C. elegans CASY-1 calsyntenin in associative learning. PloS one 4, e4880 (2009).

35. H. Ohno et al., Role of synaptic phosphatidylinositol 3-kinase in a behavioral learning response in C. elegans. Science 345, 313–317 (2014).

36. M. Toth, The other side of the coin: Hypersociability. Genes Brain Behav 18, e12512 (2019).

37. J. C. Coates, M. de Bono, Antagonistic pathways in neurons exposed to body fluid regulate social feeding in Caenorhabditis elegans. Nature 419, 925–929 (2002).

38. C. Rogers et al., Inhibition of Caenorhabditis elegans social feeding by FMRFamide-related peptide activation of NPR-1. Nat Neurosci 6, 1178–1185 (2003).

39. D. Kiser, B. Steemers, I. Branchi, J. R. Homberg, The reciprocal interaction between serotonin and social behaviour. Neurosci Biobehav Rev 36, 786–798 (2012).

40. J. Bacque-Cazenave et al., Serotonin in Animal Cognition and Behavior. Int J Mol Sci 21, (2020).

41. P. M. Whitaker-Azmitia, Behavioral and cellular consequences of increasing serotonergic activity during brain development: a role in autism? Int J Dev Neurosci 23, 75–83 (2005).

42. D. I. Zafeiriou, A. Ververi, E. Vargiami, The serotonergic system: its role in pathogenesis and early developmental treatment of autism. Curr Neuropharmacol 7, 150–157 (2009).

43. J. Golebiowska et al., Serotonin transporter deficiency alters socioemotional ultrasonic communication in rats. Sci Rep 9, 20283 (2019).

44. S. W. Flavell et al., Serotonin and the neuropeptide PDF initiate and extend opposing behavioral states in C. elegans. Cell 154, 1023–1035 (2013).

45. J. Watteyne, A. Chudinova, L. Ripoll-Sanchez, W. R. Schafer, I. Beets, Neuropeptide signaling network of Caenorhabditis elegans: from structure to behavior. Genetics 228, (2024).

46. D. Chen, K. P. Taylor, Q. Hall, J. M. Kaplan, The Neuropeptides FLP-2 and PDF-1 Act in Concert To Arouse Caenorhabditis elegans Locomotion. Genetics 204, 1151–1159 (2016).

47. A. Barrios, R. Ghosh, C. Fang, S. W. Emmons, M. M. Barr, PDF-1 neuropeptide signaling modulates a neural circuit for mate-searching behavior in C. elegans. Nat Neurosci 15, 1675–1682 (2012).

48. J. Luo, D. S. Portman, Sex-specific, pdfr-1-dependent modulation of pheromone avoidance by food abundance enables flexibility in C. elegans foraging behavior. Curr Biol 31, 4449–4461 e4444 (2021).

49. T. Janssen et al., Functional characterization of three G protein-coupled receptors for pigment dispersing factors in Caenorhabditis elegans. J Biol Chem 283, 15241–15249 (2008).

50. S. Brenner, The genetics of Caenorhabditis elegans. Genetics 77, 71–94 (1974).

51. M. Porta-de-la-Riva, L. Fontrodona, A. Villanueva, J. Ceron, Basic Caenorhabditis elegans methods: synchronization and observation. J Vis Exp, e4019 (2012).

52. C. Mello, A. Fire, DNA transformation. Methods Cell Biol 48, 451–482 (1995).

53. K. S. Ghanta, T. Ishidate, C. C. Mello, Microinjection for precision genome editing in Caenorhabditis elegans. STAR Protoc 2, 100748 (2021).

54. J. P. Concordet, M. Haeussler, CRISPOR: intuitive guide selection for CRISPR/Cas9 genome editing experiments and screens. Nucleic Acids Res 46, W242–W245 (2018).

55. M. Chalfie et al., The neural circuit for touch sensitivity in Caenorhabditis elegans. J Neurosci 5, 956–964 (1985).

56. E. K. C. Schwartz et al., Serotonin and Dopamine Mimic Glucose-Induced Reinforcement in C. elegans: Potential Role of NSM Neurons and the Serotonin Subtype 4 Receptor. Front Physiol 12, 783359 (2021).

57. S. Thapliyal, K. Babu, Pentylenetetrazole (PTZ)-induced Convulsion Assay to Determine GABAergic Defects in Caenorhabditis elegans. Bio Protoc 8, (2018).

58. B. H. Cheung, F. Arellano-Carbajal, I. Rybicki, M. de Bono, Soluble guanylate cyclases act in neurons exposed to the body fluid to promote C. elegans aggregation behavior. Curr Biol 14, 1105–1111 (2004).

59. J. Wang et al., Ciliary intrinsic mechanisms regulate dynamic ciliary extracellular vesicle release from sensory neurons. Curr Biol 34, 2756–2763 e2752 (2024).

60. C. Ding, Y. Wu, H. Dabas, M. Hammarlund, Activation of the CaMKII-Sarm1-ASK1-p38 MAP kinase pathway protects against axon degeneration caused by loss of mitochondria. Elife 11, (2022).

61. D. S. Dwyer, Crossing the Worm-Brain Barrier by Using Caenorhabditis elegans to Explore Fundamentals of Human Psychiatric Illness. Mol Neuropsychiatry 3, 170–179 (2018).

62. U. Dag et al., Dissecting the functional organization of the C. elegans serotonergic system at whole-brain scale. Cell 186, 2574–2592 e2520 (2023).

63. N. Ji et al., A neural circuit for flexible control of persistent behavioral states. Elife 10, (2021).

64. B. Chen et al., Sulfation modification of dopamine in brain regulates aggregative behavior of animals. Natl Sci Rev 9, nwab163 (2022).

65. D. S. Dwyer, P. Awatramani, R. Thakur, R. Seeni, E. J. Aamodt, Social feeding in Caenorhabditis elegans is modulated by antipsychotic drugs and calmodulin and may serve as a protophenotype for asociality. Neuropharmacology 92, 56–62 (2015).

66. J. Ben Arous, S. Laffont, D. Chatenay, Molecular and sensory basis of a food related two-state behavior in C. elegans. PloS one 4, e7584 (2009).

67. U. S. Bhat et al., FLP-15 functions through the GPCR NPR-3 to regulate local and global search behaviours in Caenorhabditis elegans. bioRxiv, 2025.2005.2002.651881 (2025).

68. S. Choi, M. Chatzigeorgiou, K. P. Taylor, W. R. Schafer, J. M. Kaplan, Analysis of NPR-1 reveals a circuit mechanism for behavioral quiescence in C. elegans. Neuron 78, 869–880 (2013).

69. K. Hong, M. Driscoll, A transmembrane domain of the putative channel subunit MEC-4 influences mechanotransduction and neurodegeneration in C. elegans. Nature 367, 470–473 (1994).

70. H. Suzuki et al., In vivo imaging of C. elegans mechanosensory neurons demonstrates a specific role for the MEC-4 channel in the process of gentle touch sensation. Neuron 39, 1005–1017 (2003).

71. T. Sugi, H. Ito, M. Nishimura, K. H. Nagai, C. elegans collectively forms dynamical networks. Nat Commun 10, 683 (2019).

72. A. Ghaddar et al., Whole-body gene expression atlas of an adult metazoan. Sci Adv 9, eadg0506 (2023).

73. M. Majeed, C. P. Liao, O. Hobert, Nervous system-wide analysis of all C. elegans cadherins reveals neuron-specific functions across multiple anatomical scales. Sci Adv 11, eads2852 (2025).

74. C. Heins et al., Collective behavior from surprise minimization. Proc Natl Acad Sci U S A 121, e2320239121 (2024).

75. D. M. Perez et al., Towering behavior and collective dispersal in Caenorhabditis nematodes. Curr Biol 35, 2980–2986 e2984 (2025).

76. C. N. Cook et al., Individual learning phenotypes drive collective behavior. Proc Natl Acad Sci U S A 117, 17949–17956 (2020).

77. A. Sehgal, E. Mignot, Genetics of sleep and sleep disorders. Cell 146, 194–207 (2011).

78. A. M. Vosko, A. Schroeder, D. H. Loh, C. S. Colwell, Vasoactive intestinal peptide and the mammalian circadian system. Gen Comp Endocrinol 152, 165–175 (2007).

79. V. Padmanabhan et al., Locomotion of C. elegans: a piecewise-harmonic curvature representation of nematode behavior. PloS one 7, e40121 (2012).

