## Supplementary material for "Neuromodulation of Swarming Behavior in *C. elegans*: Insights into the Conserved role of Calsyntenins": Combined supplemental file

**Supplementary Methods:**

**Aldicarb assay:** Aldicarb-induced paralysis assays were performed as described earlier (*1, 2*). Briefly, Nematode growth medium (NGM) plates supplemented with 1.5 mM Aldicarb [SIGMA BCBS6930V 33386-100MG] were prepared fresh from the stock solution of 100mM for each assay. Approximately 20 young adult *C. elegans* were used per replicate, with four technical replicates per genotype. The experimenter was blinded to the genotypes during scoring. The *C. elegans* were considered paralyzed if they failed to respond to gentle prodding at both the head and tail with a platinum wire on three consecutive attempts. Paralyzed individuals were removed from the plate immediately after scoring. The assay was scored every 15 minutes, and the number of paralyzed animals was recorded at each time point. Data are reported as the percentage of paralyzed *C. elegans* at 120 minutes, a time point at which maximum paralysis difference was found.

**PTZ Convulsion Assay:** This assay was performed as described in (*3*). Briefly, 30 mm NGM plates were prepared by spreading 300 µl of PTZ stock solution (100 mg/mL in absolute ethanol - SIGMA MKCR2882 P6500-25G) onto solidified 3 ml NGM to achieve a final concentration of 10 mg/ml. This was prepared on the day of the assay. Plates were dried with lids closed for 30–35 minutes. Control plates were prepared using 300 µl of ethanol. All plates were seeded with 10 µl of concentrated *E. coli* OP50 (OD ~ 6) at the center and air-dried with lids open for ~2 hours. For each assay, 10–15 synchronized, well-fed adult hermaphrodites were transferred near the bacterial lawn. After 30 minutes, animals were recorded under low-light conditions for 15 minutes using the WormLab tracking system. The *C. elegans* were scored for convulsive phenotypes: anterior convulsions (“head bobs”), wall muscle contractions (“tonic–clonic convulsions”), or full-body paralysis (“tonic convulsions”). Convulsion frequency was calculated as: % Convulsions = (Number of convulsing *C. elegans* / Total number of *C. elegans*) × 100

**Centre-Point Speed Assay:** For on-food assays, 60 mm NGM plates were prepared by freshly spotting 50–100 µl OP50 ($OD_{600}$ = 1.2–1.4) at the centre. Off-food assays were conducted on 90mm NGM plates lacking cholesterol, dried for 30 minutes under laminar air flow and equilibrated to room temperature (~22 °C) prior to use. All experiments were performed at 22–23 °C.

Individual young adult hermaphrodite *C. elegans* were transferred using halocarbon oil to an intermediate plate to remove residual bacteria and then placed on the assay plate. *C. elegans* locomotion behavior was recorded for 2 minutes using the MBF Bioscience WormLab system (Basler acA2440 camera, 1280 × 960 pixels, 7.5 fps). The pixel scale was calibrated to 8.20 µm/pixel. Each animal was tracked for 900 frames. Centre-point speed was calculated as the average displacement of the worm's midpoint coordinates per second over the entire recording. Data were extracted using WormLab software and plotted in GraphPad Prism.

**Dopamine Supplementation Assay:** Dopamine supplementation was performed in parallel with the serotonin protocol, using fresh 1 M dopamine hydrochloride stock (SIGMA BCCL0664- H8502-5G) prepared in sterile M9 buffer. *C. elegans* were pre-exposed to 40 mM dopamine for 2.5 hours on their synchronization plates (*4, 5*). Simultaneously, 60 mm NGM plates seeded with OP50 were coated with 100 µl of the dopamine solution and dried in the dark. The animals were washed and transferred to one side of the assay plate, and control animals were treated with M9 buffer only. Data shown represents the 40 mM dopamine condition.

**Figure S1:**

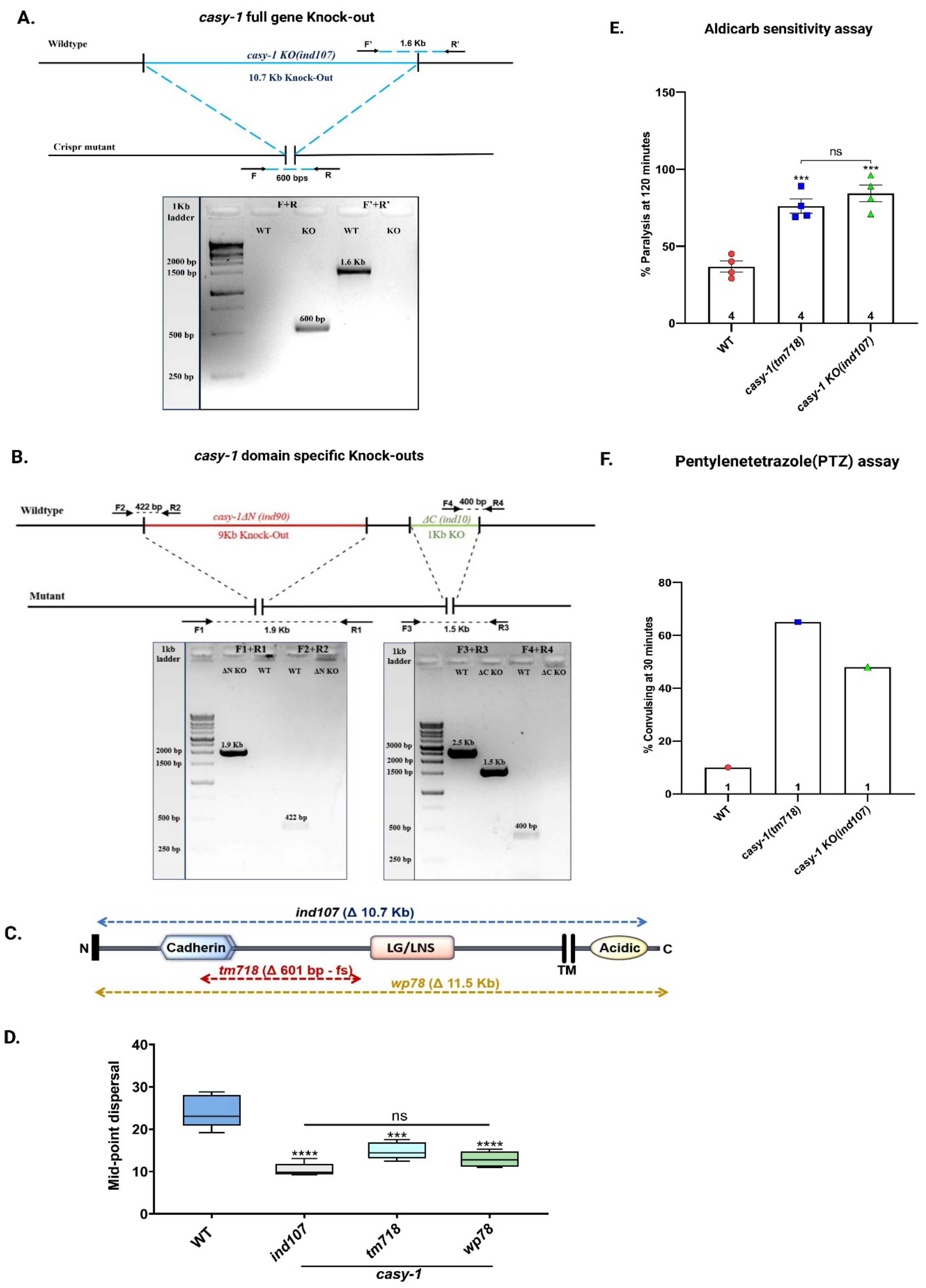
A.
casy-1 full gene Knock-out
Wildtype
casy-1 KO(ind107)
10.7 Kb Knock-Out
F'
1.6 Kb
R'
Crispr mutant
F
600 bps
R
1Kb ladder
F+R
F'+R'
WT
KO
WT
KO
2000 bp
1500 bp
1.6 Kb
600 bp
500 bp
250 bp
E.
Aldicarb sensitivity assay
% Paralysis at 120 minutes
ns
WT
casy-1(tm718)
casy-1 KO(ind107)
B.
casy-1 domain specific Knock-outs
Wildtype
422 bp
casy-1ΔN (ind90)
9Kb Knock-Out
400 bp
ΔC (ind10)
1Kb KO
Mutant
1.9 Kb
1.5 Kb
F.
Pentylenetetrazole(PTZ) assay
% Convulsing at 30 minutes
C.
ind107 (Δ 10.7 Kb)
Cadherin
LG/LNS
Acidic
TM
tm718 (Δ 601 bp - fs)
wp78 (Δ 11.5 Kb)
D.
Mid-point dispersal
ns
ind107
tm718
wp78
casy-1

**Figure S1. Generation and characterization of *casy-1* mutants and domain-specific deletions.** (A) Schematic representation of the ~10.7 kb CRISPR-mediated deletion in the genomic locus of *casy-1*(*ind107*), along with a representative genotyping gel confirming the deletion. (B) Domain-specific deletions in the *casy-1* gene: *casy-1 ΔN (ind90)* (~9 kb N-terminal deletion) and *casy-1 ΔC (ind10)* (~1 kb C-terminal deletion). A representative genotyping gel confirms the respective deletions in both mutant lines. (C) Schematic of three *casy-1* alleles: *ind107* (CRISPR KO), *tm718* (frameshift; 601 bp deletion), and *wp78* (11.5 kb deletion) with location and size of each mutation mapped relative to specific domains. Δ denotes deletion; fs indicates frameshift mutation. (D) Mid-point dispersal quantification for *casy-1* mutant alleles. *ind107* and *wp78* show reduced dispersal consistent with the swarming phenotype; *tm718* shows somewhat intermediate behaviour. (E) Aldicarb-induced paralysis assay at 120 minutes for WT, *casy-1(tm718)*, and *casy-1(ind107)* animals. (F) PTZ-induced convulsion assay showing the percentage of convulsing animals at 30 minutes across the same genotypes. This assay was performed once. Data in panels D and E represent mean ± SEM. The number of biological replicates is indicated within each bar. One-way ANOVA with Tukey's multiple comparisons test was used for statistical analysis (*** $P < 0.001$; **** $P < 0.0001$; ns is non-significant).

**Figure S2**

**A.**

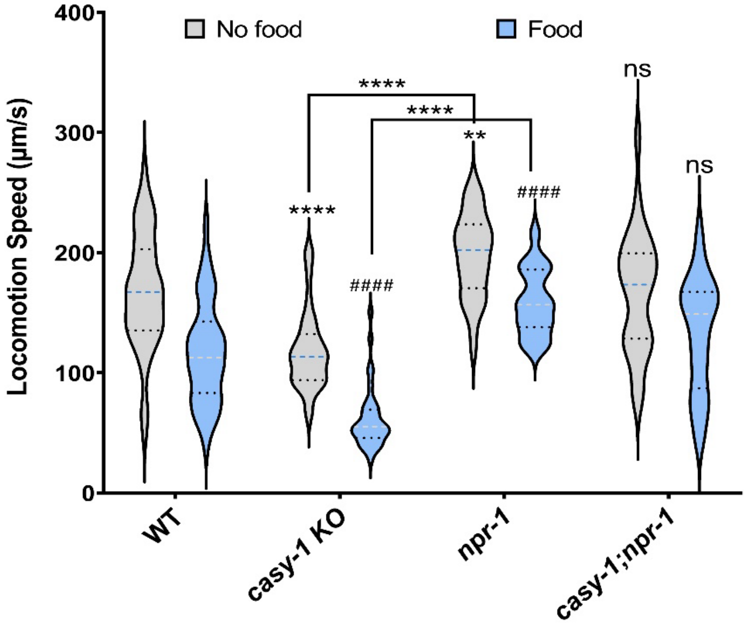
No food
Food
Locomotion Speed (µm/s)
400
300
200
100
0
****
****
**
****
####
####
ns
ns
WT
casy-1 KO
npr-1
casy-1;npr-1

**B.**

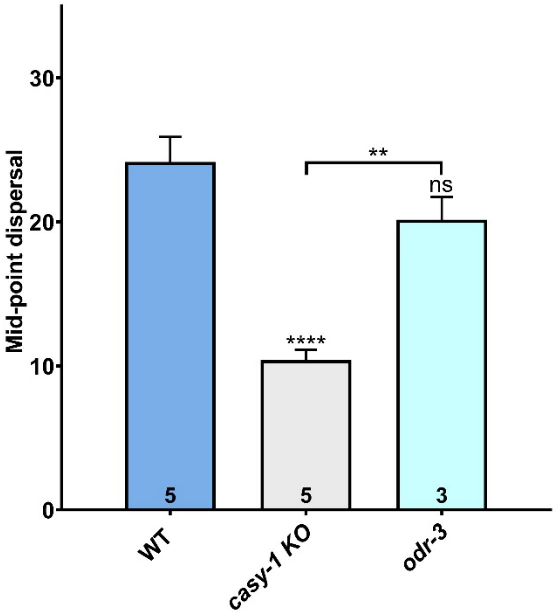
Mid-point dispersal
30
20
10
0
**
ns
****
5
5
3
WT
casy-1 KO
odr-3

**Figure S2. Locomotor speed is reduced in *casy-1* mutants, and impaired food sensation does not account for swarming.** (A) Violin plots showing center-point locomotor speed (μm/sec) for WT, *casy-1*, *npr-1*, and *casy-1; npr-1* animals under food (blue) and no-food (grey) conditions. Central dashed lines represent mean speed; outer dashed lines indicate the range of medians. *casy-1* mutants exhibit significantly reduced speed, while *npr-1* mutants display increased speed compared to WT under both conditions. *casy-1; npr-1* double mutants show no significant difference relative to WT. Statistics for WT (no food) and WT (food) are denoted by * asterisks and # hashes respectively. Sample sizes range from 35–50 animals. Two-way ANOVA (Tukey's multiple comparisons test) was used to assess significance (** $P < 0.01$, **** $P < 0.0001$; $^{\#\#\#\#}$ $P < 0.0001$, ns = not significant). (B) Mid-point dispersal for WT, *casy-1*, and *odr-3* mutants. *odr-3* mutants display WT-like dispersal, whereas *casy-1* mutants show significantly reduced dispersal. Data are represented as mean ± SEM, with replicates indicated within each bar. Statistical analysis was performed using one-way ANOVA with Tukey's multiple comparisons test. (** $P < 0.01$, **** $P < 0.0001$, ns is non-significant)

### Figure S3

A.

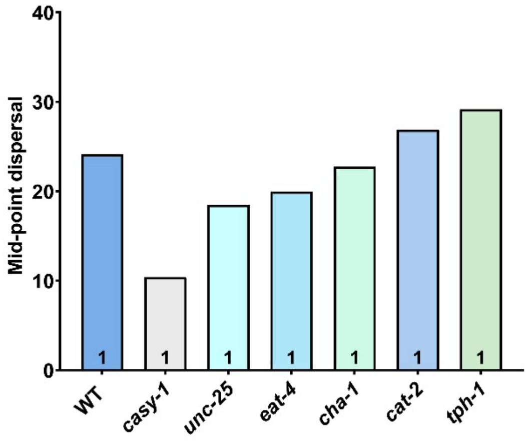


B.

| Defects | Loss-of-function Mutants |
| --- | --- |
| Neurotransmitter release | *unc-13(e51); unc-13(s69)* |
| GABA | *unc-25(tm2698)* |
| Glutamate | *eat-4(ky5)* |
| Acetylcholine | *cha-1(p503)* |
| Dopamine | *cat-2(e1112)* |
| Serotonin | *tph-1(mg280)* |

C.

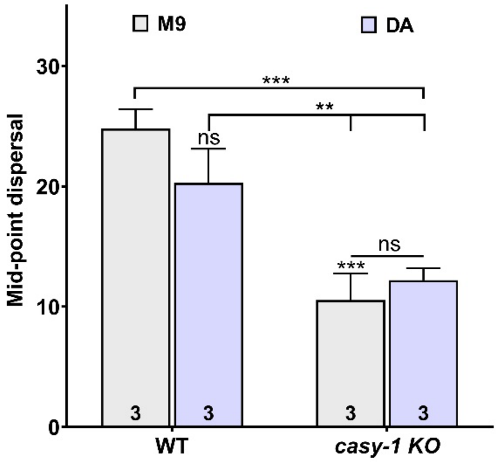


D.

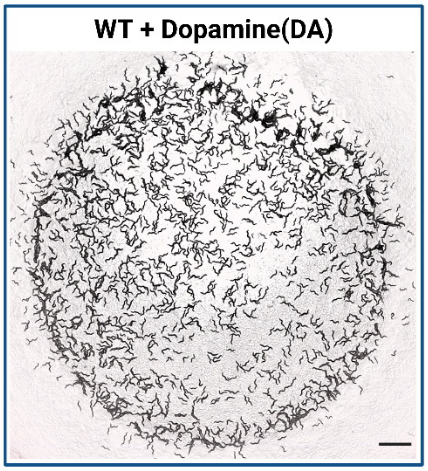

**Figure S3. Disruption of neurotransmitter signaling or dopamine supplementation does not induce swarming behavior.** (A) Mid-point dispersal for neurotransmitter pathway mutants, GABA (*unc-25*), glutamate (*eat-4*), acetylcholine (*cha-1*), dopamine (*cat-2*), and serotonin (*tph-1*), and general neurotransmitter release *unc-13* mutants. None of these mutants exhibit swarming behavior similar to *casy-1*. (B) Summary table listing the loss-of-function mutants and their corresponding neurotransmitter pathway defects. *unc-13* mutants were extremely lethargic and sick for the assay but they did not visually show any aggregation pattern. (C) Mid-point dispersal for WT and *casy-1* mutants supplemented with dopamine (DA, purple) or M9 buffer control (grey). Dopamine supplementation does not alter dispersal in either genotype. Bars represent mean ± SEM; *n* = 3 plates per group. Statistical analysis: one-way ANOVA with Tukey's post hoc test (** *P* < 0.01, *** *P* < 0.001, ns is not significant). (D) Representative image of *WT* animals supplemented with dopamine, showing a dispersed distribution across the food lawn with slight peripheral aggregation but no swarming. Scale bar: 20 mm.

**Figure S4**

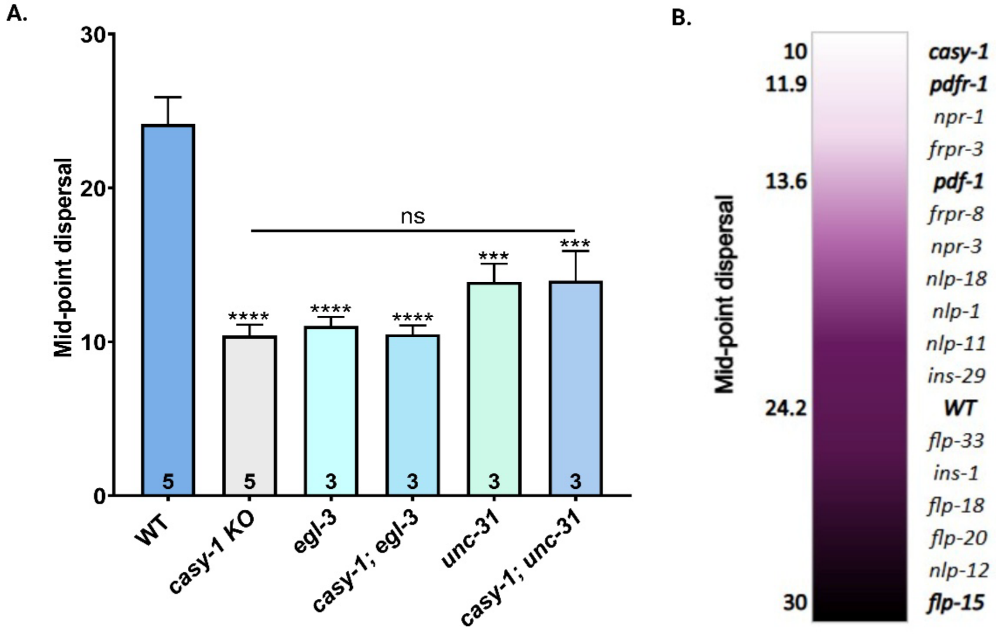

A.
Mid-point dispersal
30
20
10
0
ns
****
****
****
***
***
5
5
3
3
3
3
WT
casy-1 KO
egl-3
casy-1; egl-3
unc-31
casy-1; unc-31
B.
Mid-point dispersal
10
11.9
13.6
24.2
30
casy-1
pdfr-1
npr-1
frpr-3
pdf-1
frpr-8
npr-3
nlp-18
nlp-1
nlp-11
ins-29
WT
flp-33
ins-1
flp-18
flp-20
nlp-12
flp-15

**Figure S4. Disruption of neuropeptide signaling recapitulates swarming phenotype; sensory neuropeptide screen highlights a range of dispersal behaviors.** (A) Mid-point dispersal for *egl-3*, *unc-31*, and their respective double mutants with *casy-1*. All genotypes exhibit significantly reduced dispersal compared to WT, similar to *casy-1* single mutants, suggesting that general disruption of neuropeptide processing (*egl-3*) or release (*unc-31*) also display the swarming phenotype. Bars represent mean ± SEM; biological replicate numbers are indicated within each bar. Statistical analysis: one-way ANOVA with Tukey's multiple comparisons test (***$P < 0.001$, ****$P < 0.0001$; ns is not significant). (B) Heatmap ranking of selected neuropeptides and their receptors, primarily expressed in sensory neurons, based on their relative dispersal at midpoint (grayscale value). Lighter shades indicate lower dispersal (i.e., higher swarming), while darker shades indicate greater dispersal. *casy-1* displays the lowest dispersal (~10), while *flp-15* shows the highest (~30). Reference genotypes (*WT*, *casy-1*, *pdf-1*, and *pdfr-1*) are highlighted for comparison.

**Table S1:**

| Plasmid name | Plasmid number | Array number | Source |
|---|---|---|---|
| *Pcasy-1a::CASY-1A::T2A::mCherry* | pBAB5050 | IndEx5003 | (6) |
| *Pcasy-1a::CASY-1B::T2A::mCherry* | pBAB5150 | IndEx5007 | This Study |
| *Pcasy-1b::CASY-1A::T2A::mCherry* | pBAB5175 | IndEx5008 | This Study |
| *Pcasy-1b::CASY-1B::T2A::mCherry* | PBAB5200 | IndEx5009 | This Study |
| *Pcasy-1::mCherry* | pBAB5225 | IndEx5010 | This Study |
| *Ppdfr-1::BlaC::Sl2::GFP* | pSF154 | IndEx5012 | (7) |
| *Pcasy-1::BlaC::Sl2::GFP* | pBAB5250 | IndEx5013 | This Study |
| *Ppdf-1::BlaC::Sl2::GFP* | pBAB5275 | IndEx5014 | This Study |
| *pCFJ90-GFP* | pBAB8047 | IndEx5015 | This Study |

**Table S2:**

| Experimental Strains | Source or reference | Identifiers |
|---|---|---|
| *Wild-type Bristol strain* | CGC | N2 |
| *casy-1(ind107)* | THIS STUDY | BAB5125 |
| *casy-1(ind107); IndEx5003* | THIS STUDY | BAB5128 |
| *casy-1(tm718)* | (6) | BAB5111 |
| *casy-1(wp78)* | CGC | XE2260 |
| *casy-1(ind107); IndEx5007* | THIS STUDY | BAB5129 |
| *casy-1(ind107); IndEx5008* | THIS STUDY | BAB5130 |
| *casy-1(ind107); IndEx5009* | THIS STUDY | BAB5131 |
| *casy-1(ind90)* | THIS STUDY | BAB5126 |
| *casy-1(ind10)* | THIS STUDY | BAB5127 |
| *npr-1(ad609)* | DA609 (CGC) Outcrossed 3X | BAB5132 |
| *casy-1(ind107); npr-1(ad609)* | THIS STUDY | BAB5133 |
| *daf-22(m130)* | DR476 (CGC) Outcrossed 3X | BAB5134 |
| *daf-22(m130) casy-1(ind107)* | THIS STUDY | BAB5135 |
| *klp-6(my8); him-5(e1490)* | PT1194 (CGC) Outcrossed 3X | BAB5136 |
| *casy-1(ind107); klp-6 (my8)* | THIS STUDY | BAB5137 |
| *odr-3(n2040)* | CGC | MT4810 |
| *unc-25(tm2698)* | NBRP | Y37D8A.23 |
| *eat-4(ky5)* | CGC | MT6308 |
| *cha-1(p503)* | CGC | PR1162 |
| *tph-1(mg280)* | CGC | MT15434 |
| *cat-2(n4547)* | CGC | MT15620 |
| *bas-1(tm351)* | LC33 (CGC) Outcrossed 3X | BAB5138 |
| *casy-1(ind107); bas-1(tm351)* | THIS STUDY | BAB5139 |
| *mod-1(tm8839)* | K06C4.6 (NBRP) Outcrossed 3X | BAB5140 |
| *casy-1(ind107); mod-1(tm8839)* | THIS STUDY | BAB5141 |
| *egl-3(ok979)* | VC671 (CGC) Outcrossed 3X | BAB5142 |
| *casy-1(ind107); egl-3(ok979)* | THIS STUDY | BAB5143 |
| *unc-31(e928)* | CB928 (CGC) Outcrossed 3X | BAB5144 |
| *casy-1(ind107); unc-31(e928)* | THIS STUDY | BAB5145 |
| *pdf-1(tm1996)* | LSC27 (CGC) Outcrossed 3X | BAB6023 |
| *casy-1(ind107); pdf-1(tm1996)* | THIS STUDY | BAB5146 |
| *pdf-1(tm1996); lstls1* | Beets Lab, KU Keuven | LSC90 |
| *WT; lstls1* | THIS STUDY | BAB5147 |
| *casy-1(ind107); lstls1* | THIS STUDY | BAB5148 |
| *pdfr-1(tm12859)* | NBRP | FX34779 |
| *pdfr-1(lst34); lstEx25* | CGC | LSC60 |
| *pdfr-1(lst34)* | THIS STUDY | BAB5149 |
| *WT; lstEx25* | THIS STUDY | BAB5150 |
| *casy-1(ind107); lstEx25* | THIS STUDY | BAB5151 |
| *WT; IndEx5010; IndEx5011* | THIS STUDY | BAB5152 |
| *sls103334; IndEX5010* | THIS STUDY and BC11358 (CGC) | BAB5153 |
| *WT; IndEX5012* | THIS STUDY | BAB5154 |
| *casy-1; IndEx5012* | THIS STUDY | BAB5155 |
| *WT; IndEx5013* | THIS STUDY | BAB5156 |
| *casy-1; IndEx5013* | THIS STUDY | BAB5157 |
| *flp-15(gk1186)* | VC2504 (CGC) Outcrossed 3X | BAB6015 |
| *flp-18(gk3063)* | (6) | BAB5114 |
| *flp-20(pk1596)* | PT505 (CGC) Outcrossed 3X | BAB6020 |
| *flp-33(gk1037)* | VC2422 (CGC) Outcrossed 3X | BAB6013 |
| *nlp-11(syb530)* | PHX530 | BAB6026 |
| *nlp-12(ok335)* | RB607 (CGC) Outcrossed 3X | BAB919 |
| *nlp-18(ok1557)* | RB1372 (CGC) Outcrossed 3X | BAB6033 |
| *ins-1(nj32)* | CGC | VC581 |
| *nlp-1(ok1469)* | CGC | RB1340 |
| *ins-29(tm1922)* | NBRP | VC20254 |
| *frpr-3(ok3302)* | CGC | VC2565 |
| *frpr-8(sy1362)* | CGC | PS8486 |
| *mec-4(u253)* | CGC | TU253 |

### Table S3:

| Strains | Oligo name | Oligo sequences | Mutation |
|---|---|---|---|
| **Genotyping primers** | | | |
| *casy-1(ind107)* | NS010-F' | GACGGGTGATGGAATGAAAG | Crispr KO |
| | NS012-R' | CAGCACGCTCCTACAACAAG | |
| | NS079- F | GGTTAATCTCCACTCAATTATC | |
| | NS080- R | GTATATGTGATGTAATCAACAGGG | |
| *casy-1(tm718)* | NS010-EF | GACGGGTGATGGAATGAAAG | Deletion |
| | NS011-IR | TCAAAGCTTCTCCTCCCAGA | |
| | NS012-ER | CAGCACGCTCCTACAACAAG | |
| *casy-1(wp78)* | F | GAATAAGAATGAGAAGACCCGCTGC | Crispr KO |
| | IR | CTCCTTGCAGATTGATTATTGGCGC | [8] |
| | ER | AAGGAGTGAAAAGGACAGTATGAAGACG | |
| *casy-1 ΔC(ind10)* | NS279- F3 | GTGAATGATCTATGAGCTGAAG | Crispr KO |
| | NS280- R3 | TGAGGTCATTCAGAGAGTTG | |
| | NS223- F4 | CTCTCTGGATCCGAAATGGACCTCCCGCGTC | |
| | NS283- R4 | CTTCTCACCTGTCTCTTCTGCC | |
| *casy-1 ΔN(ind90)* | NS079- F1 | GGTTAATCTCCACTCAATTATC | Crispr KO |
| | NS080- R1 | GTATATGTGATGTAATCAACAGGG | |
| | NS281- F2 | CACATCCACATTTACTTCCTCCATTC | |
| | NS282- R2 | CCGGTTTCAGAGAGGATTGC | |
| *npr-1(ok1447)* | SS011-EF | ACCTGTCACTTTTACGCCGG | Insertion |
| | SS012-IR | TGATTTCGTTCCAGTTGAACG | |
| | SS013-ER | GAACCTTCACTTCTCCTGTG | |
| *daf-22(m130)* | NS269- Mut F | CGGTTGCTCCGATAGGATGACT | Substitution |
| | NS270- WT F | CGGTTGCTCCGATAGGATGACC | |
| | NS262- R | CCCAGTTCACCAAGGAATTCTCTC | |
| *klp-6(my8)* | F300 | TCGAAGATCTTGGCAGAGGT | Deletion |
| | R1900 | CAGATTGACGTTCGCTGAAA | [9] |
| | R1295 | TGAAACTGTTCACCGGGACT | |
| *tph-1(mg280)* | NS322- F | CGCCATCGGATATCTAAAAGAGG | Deletion |
| | NS323- Mut R | TTTGGAAACCATTCAGAACCGG | |
| | NS324- WT R | GATGCTCCAAGAGAAGCTAATCC | |
| *bas-1(tm351)* | NS333- F | GGTATCGGAAATGTGCTCGGTTC | Deletion |
| | NS334- Mut R | CAGTCACTGAAGCTCGTGCAAC | |
| | NS335- WT R | CATCGACCTGATCAACTCCACAAG | |
| *mod-1(tm8839)* | NS336- EF | TGGAACCGCTTTGAAGTTCG | Deletion |
| | NS337- IR | TCCTGAAATCACACTACTCTTGC | |
| | NS338- ER | GTTTTGGTCGCGTGATCAAC | |
| *mod-5(n822)* | NS348- F | CAGTCACCCAAACAATTTTTCCAGAC | Single-base substitution |
| | NS349- R | GTTCCGTGGGCGTCATGAGT | |
| | NS350- R | GTTCCGTGGGCGTCATGTGT | |
| *egl-3(ok979)* | NS273- F | GCATCAAGAACTCCAAATCCG | Deletion |
| | NS274- F | GTCATAGGCAACTCCGACTCC | |
| | NS275- R | GGATCCATTTTCTTGGTTGTCCC | |
| *unc-31(e928)* | NS307- F | GTATACAGAAGTGAAGAGTATGATAGC | Deletion-insertion |
| | NS308- R | CGGCTGTAAGATAGATGGAAAATT | |
| | NS310- F' | CTCCAGAAAGTTGGTAGAGTGTC | |
| *pdf-1(tm1996)* | G1 EF | CGAAATGTATTGACACTGCTGACTGC | Deletion |
| | G2 ER | GCCATGAAAATGACGGAAAATGTG | |
| | G3 IR | CTGTGCAACCTTCCTCCTTTGAAG | |
| *pdf-1(tm1996); lstls1* | NS330- F | GGAATTTTCATACTCTCCCAGCG | Deletion |
| | NS331- Mut R | GTTTTAAAAGTGGGCGTGTTGC | |
| | NS332- WT R | TGACGGAAAATGTGAAGAACGG | |
| *pdfr-1(lst34)* | NS314- F | GTCATTATTGCCACGTCAAACAC | Deletion |
| | NS315- R | GTGATCCTCGTCGAAGTTCG | |
| *mec-4(u253)* | NS386- F | GATTTGTGGAGAACATTTCGAAAC | Deletion |
| | NS387- Mut R | CTTACAAAGCAAGTTTAGCAACAAG | |
| | NS388- WT R | CATTTCCGAATTTCAGTGTGAAC | |

| Genes/Promoters | Oligo name | Oligo sequences | Vector Backbone |
|---|---|---|---|
| *Pcasy-1a* (promoter for rescue) | NS257- F | CCAAGCTATCAACTTTGTATAGAAAAGTTGCCGTACTTCCTCTGAATCGAC | pGH8 |
| | NS256- R | ATGTGGGTTTGAGTAAGAGAGCC | |
| CASY-1 A gene (for rescue) | NS245- Pcasy-1a F | CTCTTACTCAAACCCACATGGATCCATGCGAACTGCGTACTTTATTTTTGTC | |
| | NS232- Pcasy-1b F | CTTCTTTATACACTTATTTCACTGCGGATCCATGCGAACTGCGTACTTTATT TTTGTC | |
| | NS258- R | CAGACTTCCTCTGCCCTCACTAGTGACACGATAAGAACGAGCGTTC | |
| *Pcasy-1b* (promoter for rescue) | NS 260- F | GACACGATAAGAACGAGCGTTCGTTGAGATGGCG | |
| | NS261- R | GCAGTGAAATAAGTGTATAAAGAAG | |
| CASY-1 B gene (for rescue) | NS236-pCASY-1b F | CTTCTTTATACACTTATTTCACTGCGGATCCATGTTCGTGAACATTCTGG | |
| | NS237-pCASY-1a F | CTCTTACTCAAACCCACATCGGATCCATGTTCGTGAACATTCTGG | |
| | NS258- R | CAGACTTCCTCTGCCCTCACTAGTGACACGATAAGAACGAGCGTTC | |
| T2A Vector (for rescue) | NS120- F | ACTAGTGAGGGCAGAGGAAGTCTG | |
| | NS121- R | CAACTTTTCTATACAAAGTTGATAGCTTGG | |
| *Pcasy-1a* (for transcriptional reporter) | NS094- F | CTCTCTCCTGCAGGCCGTACTTCCTCTGAATCGAC | pPD49.26 |
| | NS095- R | CTCTCTGGATCCATGGTGATGTTTGGCGTAGG | |
| *Pcasy-1a* (for optogenetics) | NS389- F | GACCATGATTACGCCAAGCTTGCTCCTTTTTCGTATACCGTACTTC | pSF154 |
| | NS390- R | CATTCGAAACATACCTTTGGGTCCATGTGGGTTTGAGTAAGAGAGC | |
| *Ppdf-1* (for optogenetics) | NS398- F | GCACTGACTGGGCCGGCCTGGTCACCCAAATTAAACAAGATTAAACG | |
| | NS399- R | GGGATCCTCTAGAGGCGCGCCAGTCGAATAGAATGAATGAGCACAAC | |
| BlaC Vector | NS391- F | GGACCCAAAGGTATGTTTCGAATG | |
| | NS392- R | CAAGCTTGGCGTAATCATGGTC | |

| Gene Knock-out | PAM sequence | Oligo sequences | Donor DNA |
|---|---|---|---|
| *casy-1(ind107) 5'* | TGG | TTTGAGTAAGAGAGCCTGGT | -- |
| *casy-1(ind107) 3'* | GGG | TGTCGTTGGAGGTCTTGAGT | |
| *casy-1 ΔN(ind90) 5'* | AGG | ACGCCAAACATCACCATGCT | 5' CAAAACCAATCATTT--- |
| *casy-1 ΔN(ind90) 3'* | TGG | GTCGGACAAGGCGCGATCGC | ----CTTATTTTCATGTTA 3' |
| *casy-1 ΔC(ind10) 5'* | CGG | ACTCAAGACCTCCAACGACA | 5' GTTGTTTGCGTCG--- |
| *casy-1 ΔC(ind10) 3'* | GGG | TGTCGTTGGAGGTCTTGAGT | ---ATCTCAACGAACGCT 3' |

**References:**


1. T. R. Mahoney, S. Luo, M. L. Nonet, Analysis of synaptic transmission in Caenorhabditis elegans using an aldicarb-sensitivity assay. Nat Protoc 1, 1772-1777 (2006).
2. S. Thapliyal et al., The C-terminal of CASY-1/Calsyntenin regulates GABAergic synaptic transmission at the Caenorhabditis elegans neuromuscular junction. PLoS Genet 14, e1007263 (2018).
3. S. Thapliyal, K. Babu, Pentylenetetrazole (PTZ)-induced Convulsion Assay to Determine GABAergic Defects in Caenorhabditis elegans. Bio Protoc 8, (2018).
4. M. Baidya, M. Genovez, M. Torres, M. Y. Chao, Dopamine modulation of avoidance behavior in Caenorhabditis elegans requires the NMDA receptor NMR-1. PloS one 9, e102958 (2014).
5. P. Pandey, A. Singh, H. Kaur, A. Ghosh-Roy, K. Babu, Increased dopaminergic neurotransmission results in ethanol dependent sedative behaviors in Caenorhabditis elegans. PLoS Genet 17, e1009346 (2021).
6. N. Shahi, S. Thapliyal, K. Babu, Sensory modulation of neuropeptide signaling by CASY-1 gates cholinergic transmission at Caenorhabditis elegans neuromuscular junction. J Biosci 50, (2025).
7. S. W. Flavell et al., Serotonin and the neuropeptide PDF initiate and extend opposing behavioral states in C. elegans. Cell 154, 1023-1035 (2013).
8. C. Ding, Y. Wu, H. Dabas, M. Hammarlund, Activation of the CaMKII-Sarm1-ASK1-p38 MAP kinase pathway protects against axon degeneration caused by loss of mitochondria. Elife 11, (2022).
9. J. Wang et al., Ciliary intrinsic mechanisms regulate dynamic ciliary extracellular vesicle release from sensory neurons. Curr Biol 34, 2756-2763 e2752 (2024).
